# Anti-Inflammatory Tension-Activated Repair Patches Improve Repair After Intervertebral Disc Herniation

**DOI:** 10.1101/2022.10.22.513360

**Authors:** Ana P. Peredo, Sarah E. Gullbrand, Chet S. Friday, Briana S. Orozco, Edward D. Bonnevie, Rachel L. Hilliard, Hannah M. Zlotnick, George R. Dodge, Daeyeon Lee, Michael W. Hast, Thomas P. Schaer, Harvey E. Smith, Robert L. Mauck

**Affiliations:** Department of Bioengineering, University of Pennsylvania; Philadelphia, 19104, USA; Department of Orthopaedic Surgery, University of Pennsylvania; Philadelphia, 19104, USA; Department of Chemical and Biomolecular Engineering, University of Pennsylvania; Philadelphia, 19104, USA; Department of Clinical Studies, New Bolton Center, School of Veterinary Medicine, University of Pennsylvania; Philadelphia, PA 19348, USA; Translational Musculoskeletal Research Center, Corporal Michael J. Crescenz VA Medical Center; Philadelphia, 19104, USA

## Abstract

Conventional treatment for intervertebral disc herniation alleviates pain but does not repair the annulus fibrosus (AF), resulting in a high incidence of recurrent herniation and persistent disfunction. The lack of repair and the acute inflammation that arise after injury further compromises the disc and can result in disc-wide degeneration in the long term. To address this clinical need, we developed tension-activated repair patches (TARPs) for annular repair and the local delivery of bioactive anti-inflammatory factors. TARPs transmit physiologic strains to mechanically-activated microcapsules (MAMCs) embedded within, which activate and release encapsulated biomolecules in response to physiologic loading. Here, we demonstrate that the TARP design modulates implant biomechanical properties and regulates MAMC mechano-activation. Next, the FDA-approved anti-inflammatory molecule, interleukin 1 receptor antagonist, Anakinra, was loaded in TARPs and the effects of TARP-mediated annular repair and Anakinra delivery was evaluated in a model of annular injury in the goat cervical spine. TARPs showed robust integration with the native tissue and provided structural reinforcement at the injury site that prevented disc-wide aberrant remodeling resulting from AF detensioning. The delivery of Anakinra via TARP implantation improved the retention of disc biochemical composition through increased matrix deposition and retention at the site of annular injury. Anakinra delivery additionally attenuated the inflammatory response associated by scaffold implantation, decreasing osteolysis in adjacent vertebrae and preserving disc cellularity and matrix organization throughout the AF. These results demonstrate the translational and therapeutic potential of this novel TARP system for the treatment of intervertebral disc herniations.

**One Sentence Summary:** Tension-activated repair patches delivering bioactive anti-inflammatory factors improve healing in an in vivo goat cervical disc injury model.

## INTRODUCTION

The intervertebral disc (IVD) is a soft tissue that bridges adjacent vertebra throughout the length of the spine, providing flexibility, load transfer, and shock absorption during activities of daily living. In order to withstand the complex loads acting on the spine, the IVD has a unique configuration with regional variations in biochemical composition and tissue architecture (*1*). The disc core is a gel-like substance, termed the nucleus pulposus (NP), that has high proteoglycan and water content to withstand compressive forces. Surrounding the NP, the highly organized annulus fibrosus (AF) consists of aligned circumferential layers that enable the pressurization of the NP during compression while withstanding tension during flexion, rotation, and lateral bending (*2*). Connecting the NP/AF structure to superior and inferior vertebra are the cartilaginous endplates (CEPs). These substructures and the adjacent vertebra form the spinal motion segments which sustain and transmit physical loads during body movement.

Injuries to the disc resulting from trauma, overuse, or degeneration can cause annular ruptures and clefts (*3*). Tears that traverse the full AF thickness give way to extrusion of the NP and other disc tissue past the outer periphery of the disc. This type of injury, termed disc herniation, can result in compression of surrounding spinal nerves and chemical irritation due to elevated inflammatory cytokines, resulting in numbness and pain along the back and extremities (*4*). Although not every disc herniation results in pain, symptomatic disc herniations are prevalent, affecting 2-3% of the population (*5*). The current gold standard for surgical management of symptomatic disc herniations is microdiscectomy, in which herniated tissue is surgically excised (*6*). Although this results in nerve decompression, the underlying disc pathology is not addressed, and results in an uninhibited conduit for additional NP extrusion and a persistent compromised AF structure with impaired biomechanical function. The lack of repair, coupled with the disc’s inability to heal, makes recurrent disc herniations common, with reports of up to 25% in cases of lumbar disc herniation (*7*) and predisposes the disc to continued degenerative changes . Therefore, there is a large unmet clinical need for new repair approaches that provide annular closure and help restore disc function.

Herniated tissue causes an inflammatory response, characterized by infiltration of macrophages and an upregulation in the production of inflammatory cytokines by disc cells and leukocytes (*4, 8*). Importantly, the expression of the inflammatory cytokine interleukin 1β (IL-1β), increases with injury, aging, and degeneration (*9*). IL-1β increases the expression of matrix degrading enzymes and chemokines that recruit immune cells, establishing positive inflammatory feedback loops regulated by local concentrations of the cytokine (*10, 11*). In addition, IL-1β upregulates production of neurotrophins, signaling molecules involved in the survival, differentiation, and migration of neurons. This ingrowth of nerves may play an integral role in the onset of back pain and hyperalgesia following disc injury or degeneration (*12–14*). Hampering IL-1β−mediated catabolic signaling after annular injury might prevent further loss of extracellular matrix, decrease back pain, and improve tissue repair, and thus represents a promising therapeutic avenue for the treatment of disc herniations.

To that end, we developed a tension-activated repair patch (TARP) that acts to both provide physical closure of the ruptured disc and deliver bioactive agents to the site of injury in response to mechanical loading in the local tissue microenvironment. The base unit of the TARP is a layer of composite electrospun scaffold, designed to mimic the architecture of the AF while promoting rapid cellular infiltration (*15, 16*). These layers are assembled into the TARP via thermal stamping to include pockets containing mechanically-activated microcapsules (MAMCs) (*17, 18*) that activate in response to local deformation. We investigated the effects of TARP design on scaffold biomechanics, MAMC mechano-activation, and cellular organization. TARP stamping pattern dictated scaffold biomechanics and the transfer of local strains, impacting the mechano-activation of MAMCs under physiologic dynamic loading. To block IL-1β signaling, the FDA-approved interleukin 1 receptor antagonist, Anakinra (Sobi), was encapsulated within the MAMCs and these were integrated into the TARPs. The effects of TARP repair and TARP-mediated Anakinra delivery on the repair of annular injuries was assessed in a goat cervical disc injury model. After four weeks, TARPs remained in place at the site of implantation and integrated with the native tissue, as evidenced by robust extracellular matrix deposition throughout the TARPs. The structural support provided by TARP implantation prevented aberrant remodeling of the disc associated with AF de-tensioning post-injury, as shown by a retention of cellularity along the posterior AF and reduced necrosis and collagenous remodeling. TARP-mediated Anakinra delivery at the site of injury also improved the infiltration of repair tissue and the closure of annular injuries, which consequently preserved the normal demarcation of the NP/AF border and proteoglycan staining intensity in the NP. These results highlight the potential of this new repair system and set the stage for further translational investigation of TARPs for the repair of disc herniation in a clinical setting.

## RESULTS

### TARP Stamp Pattern Affects MAMC Patency and Scaffold Mechanical Properties

MAMCs and PCL-PEO nanofibrous scaffolds were fabricated as previously described (**Fig. 1A, Fig. S1A**)(*15, 16*). TARPs were assembled by melt stamping MAMCs between two PCL-PEO nanofibrous scaffolds (**Fig. 1B-C**). Melt stamping was performed using metal stamps that were preheated to 80°C, a temperature that, when combined with pressure, adhered the scaffold layers to form a robust multi-part assembly that could be peeled apart for visualization and analysis of encapsulated MAMCs (**Fig. S1B-C**). The combination of high temperature and applied pressure in the regions in which the stamps met the scaffold-MAMC-scaffold caused a loss of MAMC contents due to melting and/or crushing (**Fig. 1D**). This loss of MAMC contents depended on the stamp geometry. Specifically, varying the longest rhombus diagonal δ caused significant changes in MAMC patency upon stamping (**Fig. 1E**). The 3 mm δ pattern showed the greatest loss of MAMC contents upon stamping, eliciting a loss of contents in approximately 60% of MAMCs. Conversely, stamping with the 4 and 5 mm δ patterns retained the contents of a greater proportion of MAMCs, with only ∼20% and 30% of MAMCs, respectively, losing their cargo (**Fig. 1E**). Ruptured MAMCs were localized to areas that came in direct contact with the stamp (**Fig. 1D**), indicating that the loss of cargo induced by the 3 mm δ stamp occurred due to a greater contact area during stamping.

**Fig 1.**
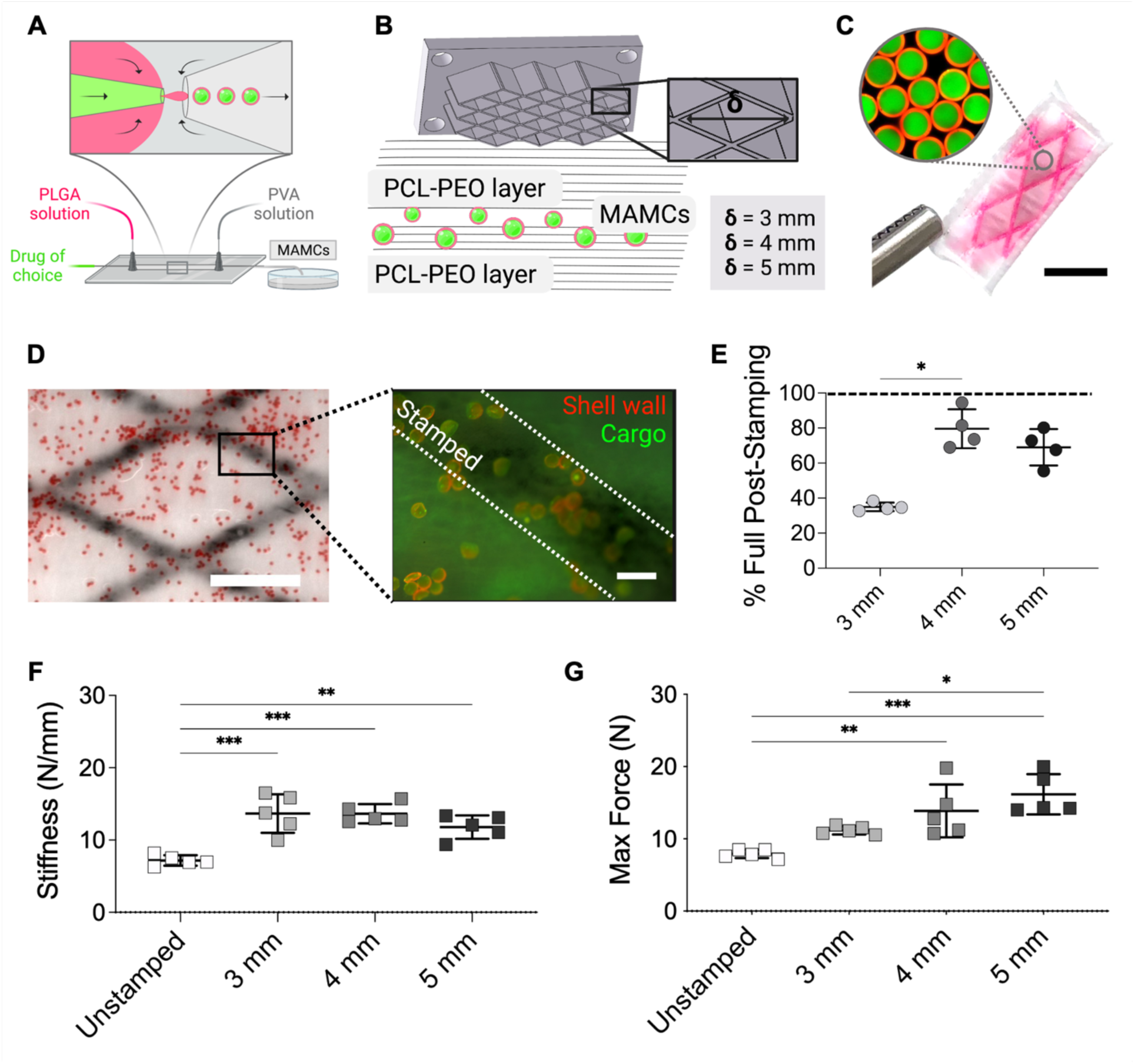
TARP fabrication with different stamp geometries regulates MAMC patency and scaffold biomechanical properties. (**A**) Schematic of MAMC fabrication using a glass capillary microfluidic device. (**B**) Schematic of TARP fabrication via melt-stamping using stamps of different rhombus patterns defined by the longest rhombus diagonal length (δ). (**C**) TARP after melt-stamping, encapsulated MAMCs seen in pink (magnified in the inset; red: MAMC shell, green: encapsulated cargo) (scale: 5 mm). (**D**) Interior of TARP imaged after melt-stamping, with MAMCs crushed/melted along the stamped region (white dashed line), verified by a loss of encapsulated cargo (green) and deformation of the MAMC shell (red) (scale:1 mm, inset scale: 100 μm). (**E**) MAMC patency, measured as % Full (mean ± s.d.), after stamping with different stamp δ (n=4 scaffolds/type). Starting %Full levels before melt stamping are indicated by the dashed line. (**F**) Stiffness and (**G**) maximum force for unstamped scaffolds or TARPs of different stamp δ (mean ± s.d.) (n=5/type). Statistical significance denoted by * p-value < 0.05, ** < 0.01, *** < 0.001.

To determine how different stamp pattern geometries impacted TARP biomechanical properties, unstamped scaffolds or TARPs were subjected to a tensile ramp to failure test, and several properties were assessed. TARP biomechanical properties changed with stamping, with differences related to the rhomboid parameter δ. Compared to unstamped scaffolds of comparable thickness, stamped TARPs had greater stiffness and maximum force (**Fig. 1F-G**), indicating that adhering the PCL-PEO layers via melt-stamping generated a composite structure with greater tensile strength than a single PCL-PEO layer. Increasing δ resulted in incremental increases in maximum force (**Fig. 1G**). TARP biomechanical properties could therefore be modulated via variations in stamp geometry.

### TARP Rhombus Dimensions Dictate MAMC Mechano-activation

To determine the role of different stamp geometries on the transfer of strain, local strains (i.e., inside each stamped rhombus) were measured using optical tracking of marked rhombus vertices during the application of uniaxial tension (**Fig. 2A**). Tension was applied along the direction of fiber alignment, which was also the direction of the longest rhombus diagonal. Comparisons between measured longitudinal local strains (ε_y_) and applied strain up to 10% revealed differences caused by variations in stamp δ (**Fig. 2B**). No significant differences were observed at physiologic outer AF strain levels (applied strain of 6%), though the mean local ε_y_ for the δ = 4 mm group was higher than that of the unstamped scaffolds (**Fig. S1D**). This trend was apparent up to the 10% applied strain. A stamp pattern of δ = 5 mm also showed ε_y_ levels above those measured on unstamped scaffolds, though to a lesser extent than for the δ = 4 mm pattern. Conversely, a δ of 3 mm hindered the transfer of strain and led to longitudinal strains less than those measured on unstamped scaffolds. Melt stamping also altered the transverse strains (ε_x_) measured across TARPs compared to unstamped scaffolds. While unstamped scaffolds had decreasing ε_x_ as ε_y_ increased, the transverse strains measured on δ = 4 mm TARPs were minimal despite an increasing ε_y_. δ = 3 and 5 mm stamped scaffolds also showed an ε_x_ that was lower than the unstamped scaffold (**Fig. S1E**). Together, these results highlighted that, compared to unstamped scaffolds, TARPs with a stamp pattern of δ = 4 mm were able to transfer greater longitudinal strains during tensile loading and that melt-stamping reduced the transfer of strain in the transverse direction.

**Fig. 2.**
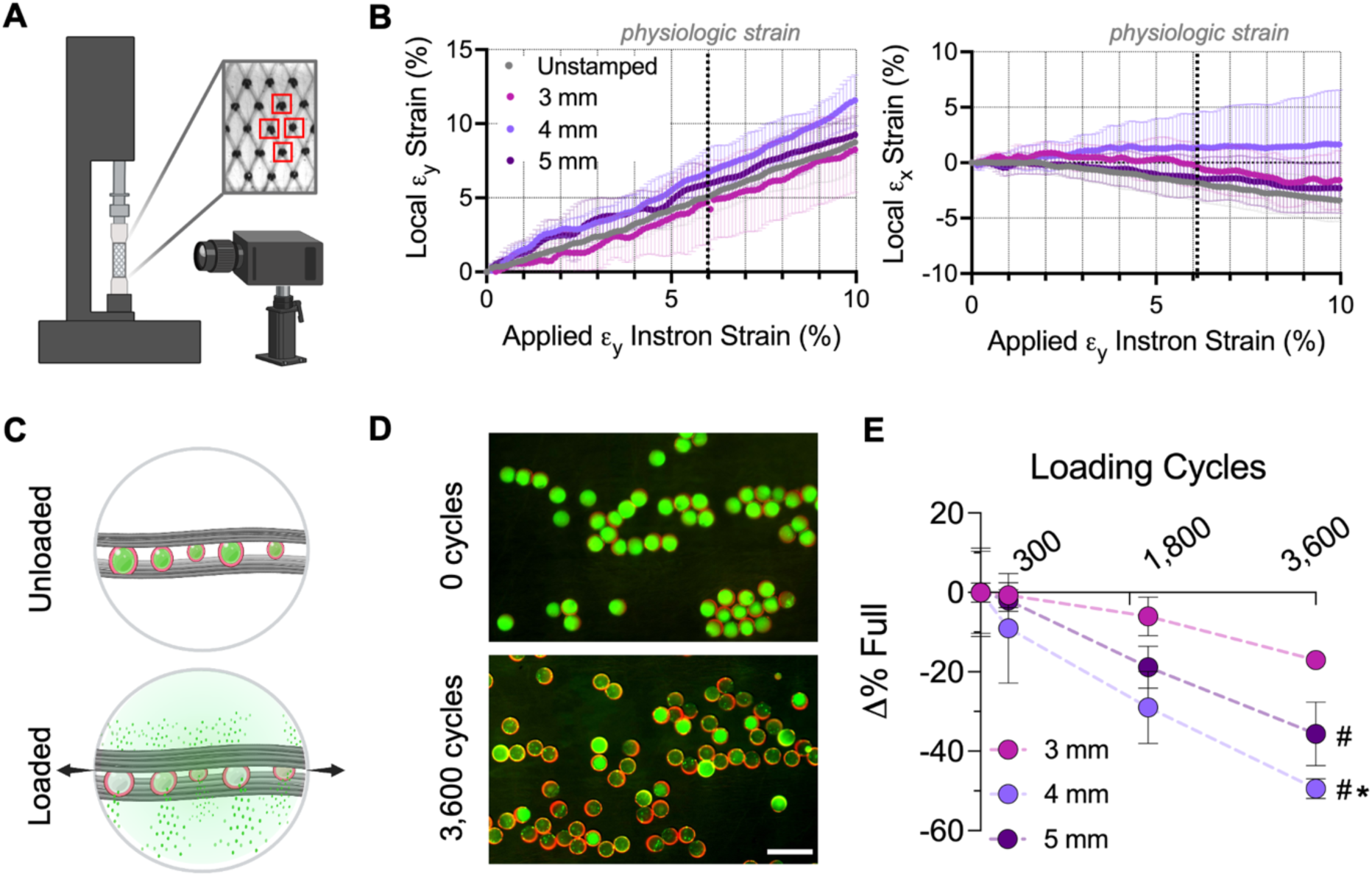
Local strain transfer dictated by stamping pattern impacts MAMC mechano-activation with dynamic physiologic tensile loading. (**A**) Schematic of optical tracking of marked rhombus vertices for local strain measurements on unstamped scaffolds and TARPs during the application of tensile ramp to failure. (**B**) Local strain in unstamped scaffolds and TARPs along the longitudinal fiber/loading direction (ε_y_) or transverse direction (ε_x_) inside marked rhombi plotted versus applied Instron strain (mean ± s.d.) (n=5/type). Physiologic outer annulus fibrosus strain (6%) highlighted by black dashed line. (**C**) Schematic of mechano-activation of MAMCs within the TARPs upon the application of dynamic tensile loading. (**D**) Representative images of MAMCs in TARP (δ = 4 mm) before the application of dynamic loading (0 cycles) or after 3,600 cycles of uniaxial tension (scale: 100 mm). Mechano-activation of MAMCs is visualized through the loss of cargo (green) from MAMCs (shell outlined in red) with TARP loading. (**E**) MAMC mechano-activation (D% Full, mean ± s.d.) for dynamic physiologic tensile loading (6% strain, 1Hz) of TARPs (n=4/type) as a function of applied loading cycles. Statistical significance denoted by # p-value < 0.05 vs. 3 mm, * p-value < 0.05 vs 4 mm.

Physiological tensile forces along the outer AF are dynamic in nature, occurring throughout daily locomotion and activity. To characterize the effects of the local strain transfer differences caused by melt stamping patterns on MAMC mechano-activation under physiologic loading (**Fig. 2C**), TARPs were assessed under dynamic tensile loading (6% strain, 1Hz) using a custom motorized uniaxial bioreactor. The roles of stamp pattern and loading cycles on the mechano-activation of MAMCs were determined by measuring % Full MAMCs with and without loading (**Fig. 2D**). Increasing cycles of loading led to incremental mechano-activation of MAMCs, irrespective of stamping pattern. After 300 loading cycles, there were no differences in % Full compared to unloaded TARPs, regardless of stamp δ (**Fig. 2E**). However, after 1,800 and 3,600 cycles of loading, significant differences were observed between different TARP designs (**Fig. 2D-E**). Specifically, larger stamp δ patterns caused greater mechano-activation, with δ = 4 and 5 mm showing increased levels of MAMC mechano-activation compared to the δ = 3 mm TARPs. Interestingly, the 4 mm δ TARPs showed the greatest mechano-activation compared to other TARP stamp patterns (**Fig. 2D**). After 3,600 cycles, ∼50% of MAMCs mechano-activated in the δ = 4 mm TARPs while ∼38% and 18% of MAMCs ruptured in the 5 mm and 3 mm δ TARPs. The increased mechano-activation of MAMCs for the 4 mm δ stamp design coincided with the greatest strain transfer measured in the longitudinal direction (and least transverse strain), suggesting that the strain transfer regulated by stamp design directly impacted the mechano-activation of TARP-encapsulated MAMCs.

The process of melt stamping generates curvature emanating from the points of melt stamping in which pressure and increased temperature anneal the scaffold layers together (**Fig. S2A**). To understand how this curvature affected the alignment within the scaffold layers, fibers were visualized before and after melt stamping and alignment was measured. Melt stamping increased fiber alignment, observed through a decrease in the fiber offset from 90° (**Fig. S2B-C**) after stamping. Larger rhombus δ (4 and 5 mm) patterns significantly increased fiber alignment compared to unstamped scaffolds. This was accompanied by a reduction in fiber angle dispersity – a measure of the heterogeneity or spread in fiber alignment – regardless of stamp δ (**Fig. S2B-C**). These results indicate that melt stamping increased fiber alignment and reduced fiber dispersity, with greater fiber realignment observed in TARPs created with larger stamp δ.

Nanofibrous scaffolds present topographical cues through their nano-architecture and organization that affect cellular sensing, and ultimately, cellular morphology. To assess the effect of changes in fiber alignment on cellular morphology, human AF cells were cultured on unstamped scaffolds or on TARPs of different stamp δ. After 6 days of culture on each material, cells were stained and visualized. As expected, cells showed increased alignment with melt stamping, which corroborated the increased fiber alignment (**Fig. S2D**). This increase in cellular organization was accompanied by an increased cell aspect ratio on melt stamped TARPs, with cells demonstrating highly extended cell bodies along the fibers (**Fig. S2E-F, Supplemental Movie 1**). Melt stamping therefore causes a structural reorganization of nanofibers, resulting in increases in fiber and cellular alignment with stamping, impacting cellular interactions.

### MAMC-mediated Delivery of Anakinra Attenuates Catabolic Gene Expression in AF Cells

The FDA-approved interleukin 1 receptor antagonist, Anakinra, has potent IL-1β inhibitory effects, including when delivered from MAMCs (*18, 19*). Although this previous study verified that MAMC fabrication did not affect bioactivity, the effects of Anakinra, particularly when delivered from MAMCs, was not investigated on AF cells. To verify that MAMC-mediated Anakinra delivery effectively inhibited IL-1β signaling, human AF cells were cultured in the presence of IL-1β and then treated with Anakinra delivered directly into the media (soluble) or extracted from MAMC contents before media supplementation (**Fig. S3A-B**). Gene expression analysis for the catabolic genes MMP3 and LCN2 demonstrated a significant upregulation when the cells were exposed to IL-1β without the addition of Anakinra. This upregulation was drastically reduced in a dose-dependent manner with the addition of Anakinra, both in the soluble and MAMC-extracted form (**Fig. S3C**). Assessments of cell viability verified that Anakinra concentrations of up to 1,000 ng/mL did not have a detrimental effect on AF cell survival (**Fig. S3D**). Together, these results demonstrated the potential of Anakinra provision to attenuate IL-1β-mediated catabolic signaling in AF cells.

### TARP-mediated Delivery of Anakinra in a Goat Cervical Spine Disc Herniation Model

To determine if TARPs can aid in the repair of annular tears with or without the delivery of an anti-inflammatory molecule (IL-1ra, Anakinra), empty TARPs and Anakinra-loaded TARPs were implanted *in vivo* in a large animal cervical disc annular injury model for up to 4 weeks. Due to the increased retention of MAMC contents post-stamping, higher strain transfer, and MAMC mechano-activation under dynamic tensile loading observed for the 4 mm δ TARPs, this stamp geometry was chosen for this *in vivo* study. TARPs were assembled with dimensions previously determined to fit within the goat cervical disc space, loaded with MAMCs containing a BSA model drug (E-scaffold) or Anakinra (A-scaffold). A total of n=8 goats underwent surgery, where the C2-3 and C3-4 cervical discs received annular injury followed by TARP repair of one injured level per animal (**Fig. 3A-B**). Adjacent levels were used as uninjured controls. The annular injury consisted of a partial thickness X-shaped laceration on the anterior AF followed by a full thickness needle puncture, mimicking the full-thickness tears that give way to NP leakage and subsequent disc herniation (**Fig. 3B-C**). Levels that underwent TARP repair received either E-scaffolds or A-scaffolds, which were sutured in place over the annular injuries (**Fig. 3D-E**). There were no surgical complications arising from the disc injuries or the TARP implantation, and all animals recovered uneventfully until the study end date.

**Fig 3.**
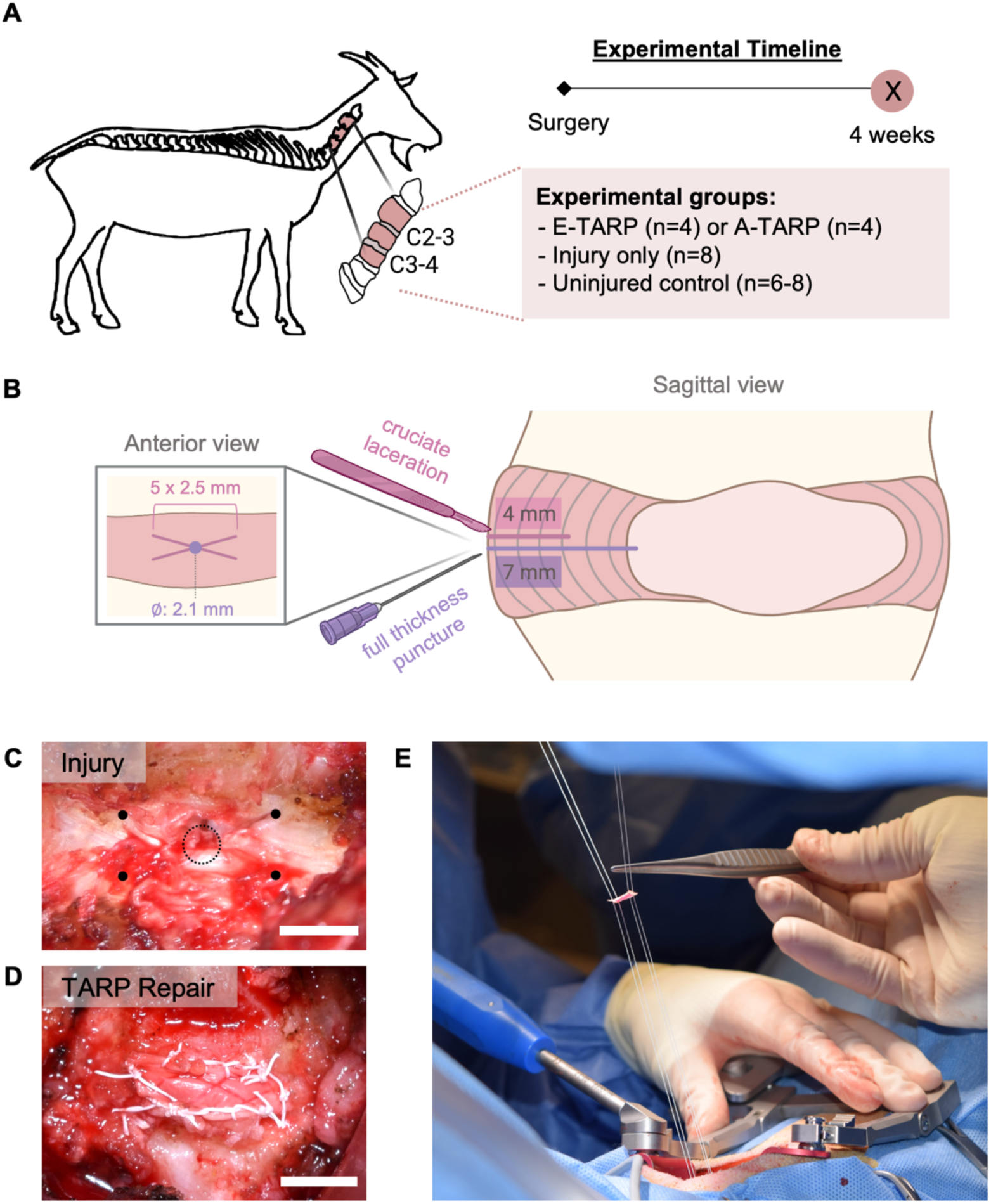
In vivo repair via TARP implantation in a goat cervical disc annular injury model. (**A**) Study design for the in vivo study using a goat cervical disc annular injury model. (**B**) Depiction of the annular injuries induced on the anterior aspect of the intervertebral disc. (**C**) Intra-operative photograph of the anterior cervical disc after annular rupture, with vertices of the X-laceration demarcated by black dots and the needle puncture area demarcated with a dashed black circle (scale: 5 mm). (**D**) Intra-operative photograph of the cervical disc after repair with a TARP, sutured across all edges to the native tissue (scale: 5 mm). (**E**) Photograph showing pre-threading of sutures through the corners of the cruciate annulotomy and the TARP being advanced along the sutures in order to secure each corner at the X-laceration and ensure central placement across the injury.

TARPs remained securely attached at the site of implantation after 4 weeks. Integration of the TARPs with the native tissue was observed through H&E staining of histological sections (**Fig. 4**), which showed robust deposition of extracellular matrix next to and throughout the expanse of each TARP (**Fig. 4C-F**). MAMC outlines were still visible after 4 weeks (**Fig. 4F**). MicroCT analysis revealed that E-Scaffold-treated levels had osteolysis along the anterior aspect of the adjacent vertebras while discs that only received injury did not show signs of bone loss (**Fig. S4A**). Interestingly, Anakinra-loaded TARPs did not elicit significant osteolysis and were comparable to uninjured control levels (**Fig. S4B**), suggesting that Anakinra delivery attenuated the inflammation-mediated loss of bone caused by TARP implantation. To determine if other changes in bone microarchitecture occurred as a result of injury or TARP repair, the cranial and caudal bony endplates adjacent to the discs were contoured and evaluated by microCT. Bone morphometric analyses revealed no differences between groups in bone volume/total volume, trabecular number, and trabecular spacing (**Fig. S4C-F**).

**Fig. 4.**
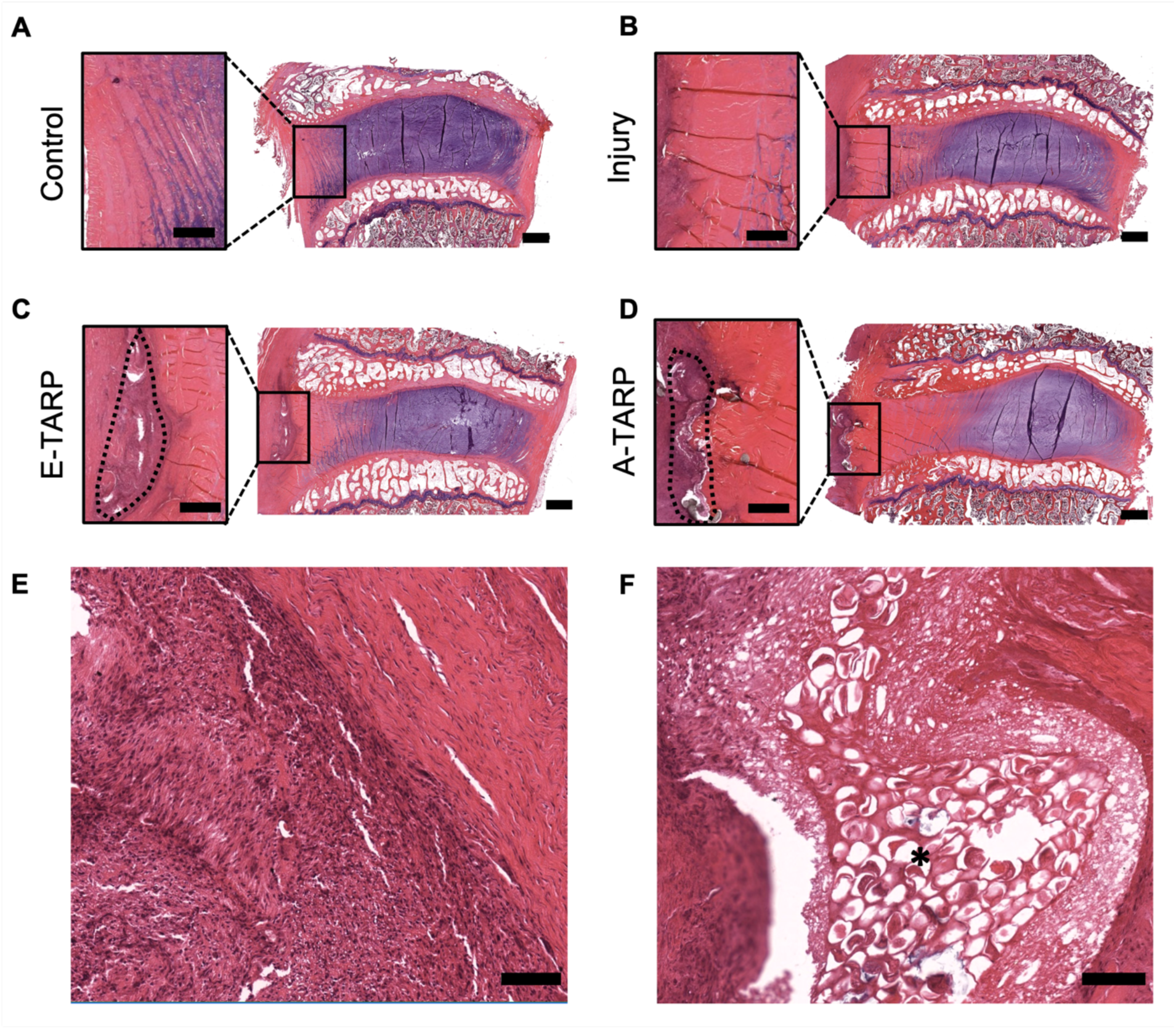
TARPs show robust integration with native tissue. **(A)** Representative H&E-stained sagittal section and zoom in of anterior AF for cervical discs in the uninjured control, (**B**) injury, (**C**) E-scaffold-treated, and (**D**) A-scaffold-treated groups (scale: 2 mm, inset scale: 1 mm). (**E**) Representative H&E-stained section for E-scaffold and (**F**) A-scaffold-treated discs, showing cellular infiltration of the empty and Anakinra-loaded TARPs with robust extracellular matrix deposition throughout the scaffold and integration with host tissue (scale: 100 μm). MAMC outlines (black asterisk) are apparent inside Anakinra-loaded TARP.

T2-weighted magnetic resonance imaging (MRI) is commonly used to assess disc health and NP T2 relaxation times correlate with disc hydration and proteoglycan content (*20, 21*). Average T2 maps for each group revealed differences in AF T2 and NP/AF border morphology (**Fig. 5A-B**, **Fig. S5A**). In uninjured discs, NP T2 signal is centrally condensed and surrounded by a lower AF T2 signal. With injury, this pattern spread laterally towards the anterior and posterior AF (**Fig. 5B**, **Fig. S5A**). This resulted in a significant increase in AF T2 (**Fig. S5B**), measured at the anterior AF, while NP T2 was unchanged (**Fig. S5C**). Repair of injured discs with TARPs caused a greater retention of the NP/AF border compared to injured discs (**Fig. 5A-B**), suggestive of a faster closure of the injury tract that may have prevented NP leakage or spreading. While repair with E-scaffold increased the anterior AF T2, annular repair with the A-scaffold did not affect AF T2 (**Fig. 5C**). This suggests that Anakinra delivery may have attenuated an inflammatory response induced by E-scaffold implantation, consistent with the reduced osteolysis findings noted above. There were no differences in NP T2 across all treatment groups (**Fig. 5D**) despite an apparent anterior shift of the NP with injury. Overall, this MRI data suggested that the TARPs improved the repair of the AF injury tract and prevented anterior spreading of the NP, and that Anakinra delivery attenuated inflammatory remodeling at the site of TARP implantation.

**Fig. 5.**
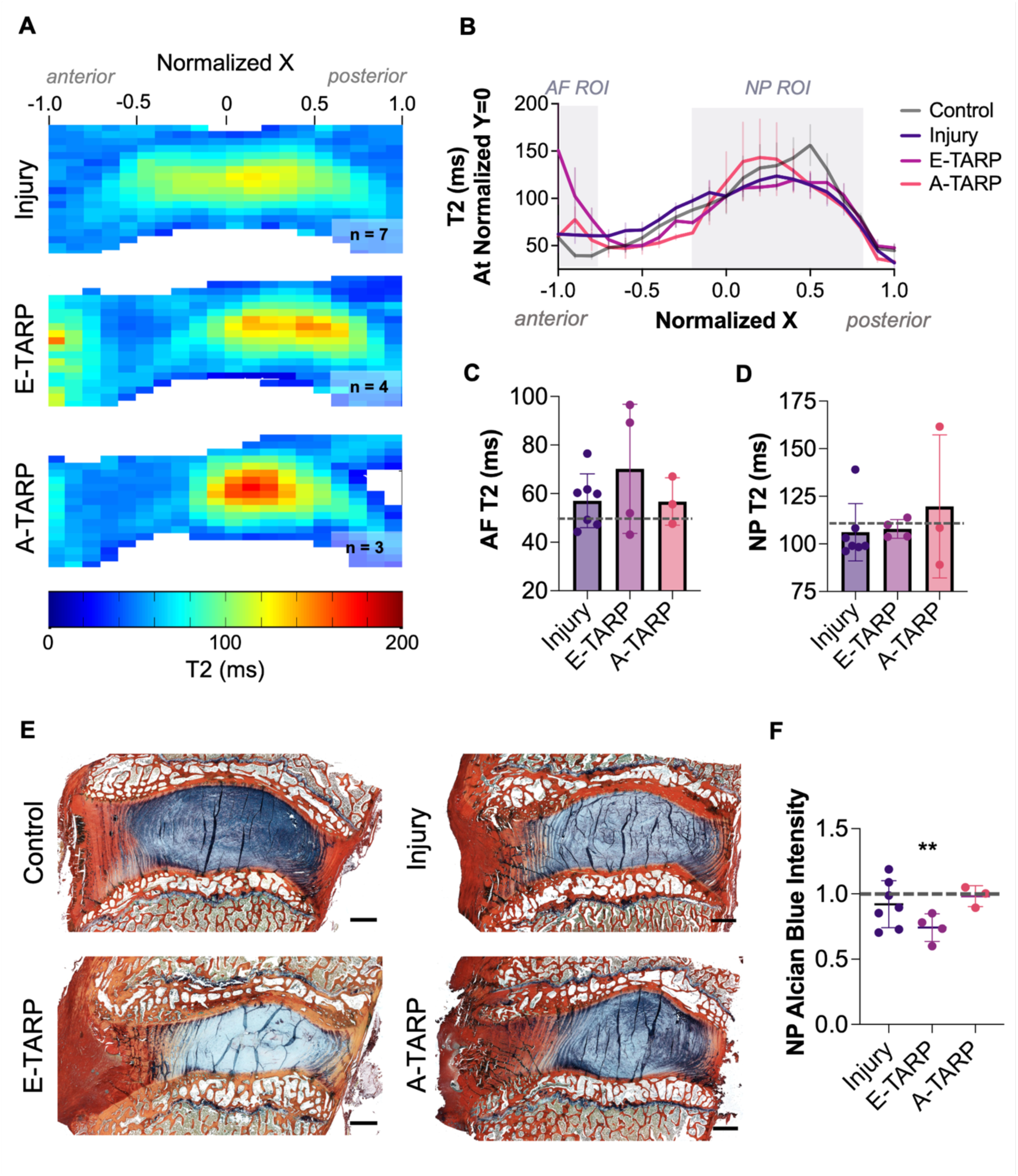
Effects of TARP repair and Anakinra delivery on disc T2 signal and biochemical composition. (**A**) Average sagittal T2 maps, with dimensions normalized to account for disc curvature and anatomical differences, for levels with injury (n=7), repair with empty TARP (E-Scaffold, n=4), and repair with Anakinra-loaded TARP (A-Scaffold, n=3). (**B**) Average T2 (mean ± SEM) at central Y axis (y = 0) of normalized T2 maps along the normalized X dimension. Anterior AF and NP regions measured through ROIs, quantified in C-D appear shaded in. (**C**) AF and (**D**) NP T2 measurements (mean ± s.d.) for rectangular and circular regions of interest, respectively, for treated discs compared to uninjured controls (dashed line: control mean). (**E**) Alcian Blue/Picrosirius Red-stained sagittal sections for discs in each group with the highest NP Alcian Blue staining intensity (scale: 2 mm). (**F**) NP Alcian Blue intensity measurements (mean ± s.d.) for sections from treated groups compared to uninjured control discs (dashed line). Statistical significance compared to control denoted by ** p-value < 0.01.

To determine if the differences in T2 relaxation times were caused by changes in biochemical composition, sagittal histological sections were co-stained with Alcian Blue and Picrosirius Red to visualize proteoglycans and collagens, respectively. Stark differences in NP proteoglycan staining intensity were observed between groups and this was verified through quantification of Alcian Blue staining intensity in the NP region of interest (**Fig. 5E-F, Fig. S6**). As expected, the highest proteoglycan staining intensity was measured in uninjured control discs (**Fig. 5F**). Injury caused a slight reduction in NP proteoglycan staining intensity compared to controls, albeit not statistically significant. E-Scaffold-treated discs showed the greatest reduction in NP proteoglycan staining intensity and was significantly lower than uninjured controls (**Fig. 5F**). Interestingly, TARP-mediated Anakinra delivery prevented this loss of NP proteoglycans, resulting in proteoglycan staining comparable to that of uninjured discs (**Fig. 5F**). Intense NP proteoglycan staining was apparent across all A-Scaffold discs analyzed when compared to the best, median, and worst specimens for each group, selected based on NP Alcian Blue intensity (**Fig. S6**). This suggested that Anakinra delivery via TARP implantation helped retain disc biochemical composition after injury, in accordance with the MRI T2 data.

The large differences in proteoglycan staining intensity between E-Scaffold and A-Scaffold-treated discs suggested differences in healing response. To determine how these differences arose, we examined the injury location more closely. Sagittal histological sections were stained with hematoxylin and eosin (H&E) for visualization of cellular morphology and Mallory Heidenhain stain for inspection of scar infiltration and collagenous remodeling. Close inspection of the anterior AF revealed a loss of cellularity near the injury tract irrespective of treatment (**Fig. 6A-D**), suggesting that tissue injury induced necrosis and apoptosis near the site of insult, consistent with previous reports (*22–24*). Mallory Heidenhain staining revealed differences in scar tissue infiltration at the anterior AF (**Fig. 6A-D**), verified through quantification of scar tissue infiltration area (**Fig. 6E**). Granulation tissue composed mainly of collagen was stained with a dark purple color, showing infiltration of scar tissue along the outer third of the anterior AF for injury and E-Scaffold groups (**Fig. 6A-E**). In contrast, injury repair with Anakinra-loaded TARPs (A-Scaffold) resulted in greater infiltration and closure of the injury tract along the anterior AF (**Fig. 6D-E**). The greater infiltration of scar tissue and consequent closure of the annular injury may have prevented shifting or loss of NP material, resulting in greater retention of the NP/AF boundary (**Fig. 5A**) and NP proteoglycan content (**Fig. 5E-F**). This was supported by the differences in scar tissue infiltration and retention of the NP/AF border that was apparent in the H&E and Mallory Heidenhain-stained sections that represented best, median, and worst sections chosen based on NP Alcian Blue intensity (**Fig. S7, Fig S8**).

**Fig. 6.**
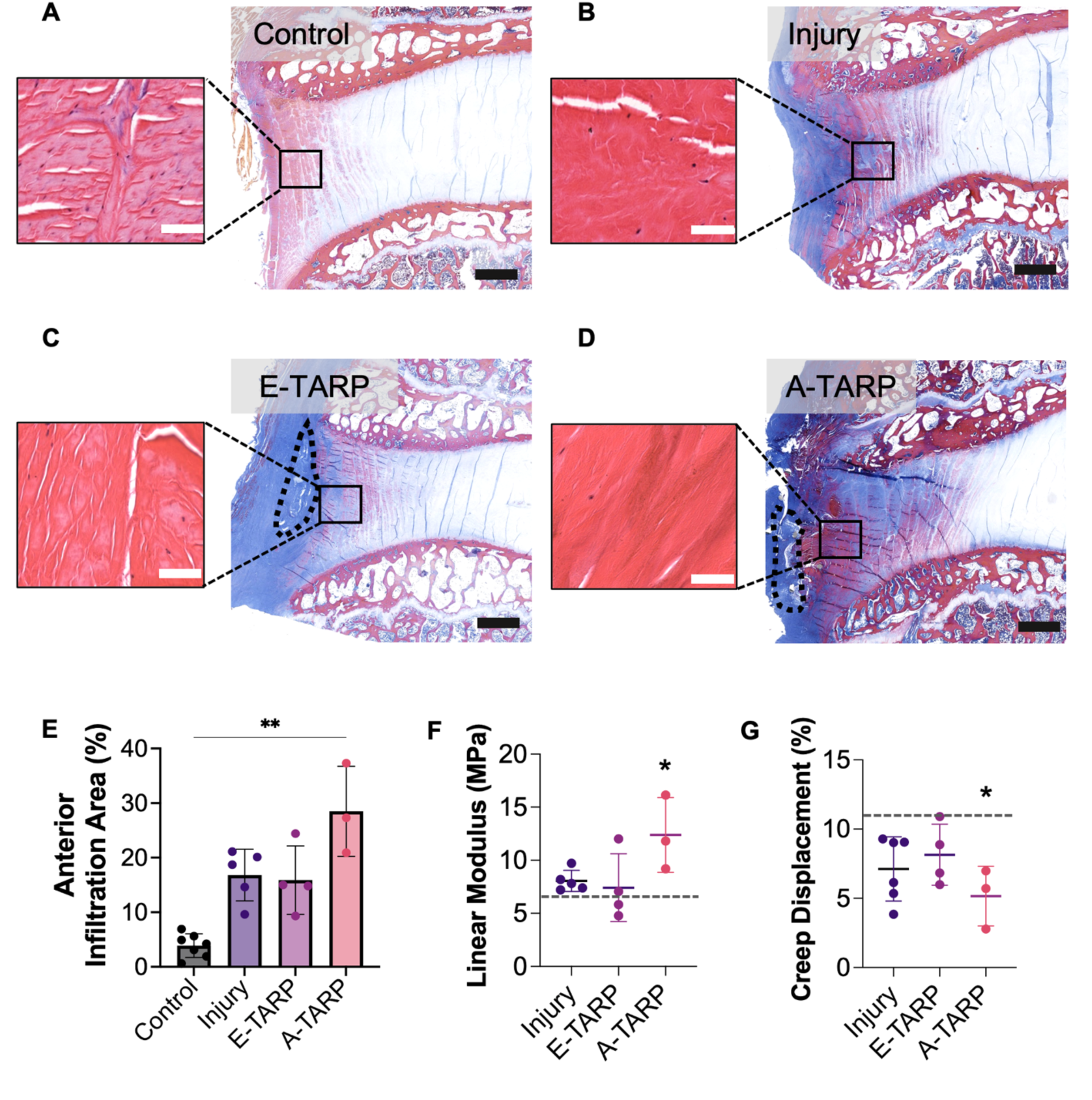
Collagenous infiltration through the injury tract increases with Anakinra delivery. (**A**) Representative Mallory Heidenhain-stained and zoomed-in H&E-stained sagittal section of anterior AF for control, (**B**) injury, (**C**) E-Scaffold, and (**D**) A-Scaffold treated discs (scale: 2 mm; TARPs are outlined with black dashed line; inset scale: 50 μm). (**E**) Anterior AF infiltration area of collagenous scar/remodeling, stained in purple, measured as percentage of total disc area from Mallory Heindenhain-stained sections (mean ± s.d.). Statistical significance denoted by ** p-value < 0.01. (**F**) Linear modulus and (**G**) creep displacement for motion segments in each group compared to uninjured controls (dashed line: control mean). Statistical significance denoted by * p-value < 0.05 vs uninjured control.

The extent of anterior scar infiltration affected disc biomechanics, determined through motion segment testing under cyclic compression and compressive creep at physiologic loads. Injured and E-Scaffold-treated discs, which had similar levels of anterior scar infiltration, showed comparable degrees of disc stiffening (**Fig. 6E, Fig. S5D-G, Fig. S9**). The levels repaired with A-Scaffolds showed greater stiffening with significant differences in linear modulus and creep displacement, compared to uninjured controls (**Fig. 6E, Fig. S5D-G**). This is suggestive of stiffening of the disc after 4 weeks of TARP-mediated Anakinra delivery and disc repair.

Annular lesions and herniation are related to degeneration of the surrounding tissue as a consequence of detensioning of the AF that causes disc-wide aberrant remodeling (*24*). For this reason, the posterior AF was inspected for signs of catabolic changes. Close analysis of H&E-stained sections along the posterior AF revealed dramatic differences in cellularity among groups (**Fig. 7A**). Quantification of cell nuclei across three regions of interest along the length of the posterior AF revealed a marked loss of cells, characteristic of tissue necrosis, along the posterior AF of injured discs that did not receive repair (**Fig. 7B**). In contrast, both E-Scaffold and A-Scaffold-treated discs retained cellularity levels comparable to uninjured controls, suggesting that TARP delivery provided mechanical reinforcement of the injured AF that prevented degenerative remodeling across the disc (**Fig. 7B**). Close inspection of Mallory Heidenhain-stained histological sections revealed differences in collagenous remodeling along the posterior region of the AF. Quantification of this posterior remodeling revealed an increase in collagenous remodeling with injury (**Fig. 7C**). Interestingly, TARP implantation, regardless of MAMC content, maintained posterior AF remodeling at uninjured control levels (**Fig. 7F**), in keeping with the posterior AF cellularity and necrosis findings. Together these results highlight that TARP implantation provides reinforcement of the AF, preventing disc-wide remodeling and tissue necrosis after injury.

**Fig. 7.**
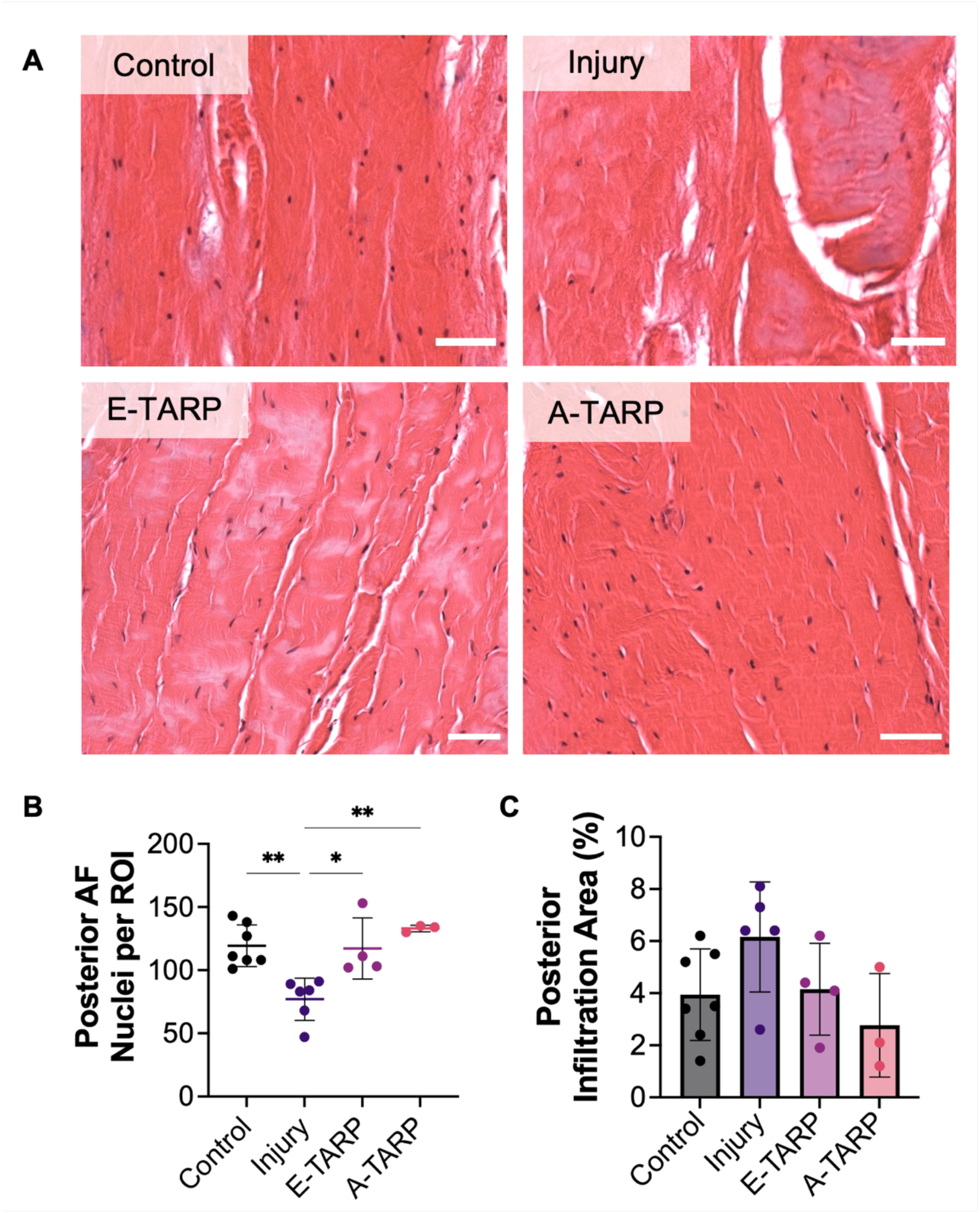
TARP implantation prevents disc-wide aberrant remodeling and tissue necrosis. (**A**) Representative H&E-stained sagittal histological sections, zoomed in on the posterior AF for control, injury, E-Scaffold, and A-Scaffold treated discs (scale: 50 μm). (**B**) Average number of nuclei per ROI in the posterior AF, measured from 3 ROIs along the outer AF of each section, using H&E-stained sagittal sections (mean ± s.d.). (**C**) Posterior AF infiltration area of collagenous scar/remodeling, stained in purple, measured as a percentage of total disc area using Mallory Heindenhain-stained sagittal sections (mean ± s.d.). Statistical significance denoted by * p-value < 0.05, ** < 0.01.

## DISCUSSION

The repair of AF injuries via an annular patch that restores the compromised disc structure and simultaneously delivers factors to aid in healing holds promise as a treatment approach for herniated intervertebral discs. There are currently no FDA-approved devices for the repair of the disc structure following herniation injury or the prolonged provision of biologic agents to the AF. A successful repair device must restore the combined function of all disc substructures to ensure its effective functioning as a single load-bearing unit. The device must also cause minimal damage to the native tissue upon implantation and must also provide reinforcement at the injury site to prevent degenerative changes that result from AF detensioning. Biologic supplementation of the avascular disc to attenuate inflammation at the injury site can further aid in tissue regeneration by limiting degradation of disc ECM and cellular apoptosis. For clinical translation, the repair device must demonstrate successful mechanical and biological support of IVDs of comparable anatomy, size, and mechanics to human discs. These challenges have impeded the translation of effective repair approaches to the clinic.

In this study, we developed and evaluated the tunable design of TARPs for closure of annular lesions and provision of anti-inflammatory molecules locally to the injury site. Through modifications of the TARP stamping pattern, biomechanical properties and rate of MAMC mechano-activation were varied, providing flexibility in the design of a repair device with a set of desired characteristics. The MAMC drug delivery system offers added tunability over drug provision, with the ability to choose MAMC rate of degradation and sensitivity to mechanical loading through the choice of PLGA employed and modification of microcapsule dimensions during fabrication (*17, 18*). A single population or a combination of MAMC populations with different characteristics can therefore be used to provide delivery of a range of molecules at different times post-implantation. In this study, MAMCs were used to deliver the FDA-approved small molecule Anakinra, which effectively inhibited the upregulation of catabolic genes in AF cells cultured with IL-1β, demonstrating the promise of this therapeutic target for the prevention of matrix degeneration post-injury.

Building on these promising in vitro results, we translated empty or Anakinra-loaded TARPs for the repair of annular injuries in the goat cervical spine for up to 4 weeks. The goat cervical spine is an attractive preclinical model due to its semi-upright stature and its approximation of human cervical spine dimensions (*25*). Our results demonstrated robust retention, infiltration, and integration of the TARP implants with the native tissue. Implantation of TARPs alone, acting as a physical barrier, was sufficient to prevent tissue-wide degenerative changes through the provision of structural reinforcement of the compromised AF. TARP-mediated Anakinra provision at the repair site further improved the retention of the distinct sub-compartments of the disc, maintaining the NP/AF boundary by preventing NP displacement. Anakinra delivery also attenuated the inflammatory response associated with device implantation, which often can lead to device failure and further catabolic changes at the implantation site.

Due to the dense composition of the fibrocartilaginous AF, attachment of annular repair devices remains a challenge. The only FDA-approved device intended to physically block recurrent herniation is the Barricaid Annular Closure device (Intrinsic Therapeutics, Inc., Woburn, MA) -a bone-anchored implant with a polymer fabric end that is placed adjacent to the AF herniation (*26, 27*). Although this device avoids damaging the AF during implantation, the attachment sites at the adjacent vertebras, where metal screws are inserted, have shown significant osteolysis in preclinical investigations (*28*). To avoid this issue, most work in the field thus far has been focused on the development of gel-based sealants or adhesives to seal AF lesions (*29*). Although adhesives provide ease of delivery via direct injection into the injury site, several have shown inadequate retention of disc mechanical function and have failed to prevent degenerative changes, possibly due to the lack of infiltration and deposition of matrix by endogenous cells (*30–32*). In this study, we demonstrated successful attachment of the TARPs to the AF using micro sutures, which enabled the retention of the TARPs at the implantation site after 4 weeks of unrestricted physical activity. The retention of the scaffold at the injury site permitted the cellular infiltration of TARPs by native cells that deposited dense matrix throughout the entirety of the scaffolds, which in turn ensured robust integration with the surrounding tissue. Although an effective method for scaffold attachment, manual suturing of the TARPs presents challenges including a lengthy procedure time and, at times, limited access caused by the size and curvature of the spine. An alternative method that partially automates the suturing and provides greater ease of implantation could be beneficial for future clinical translation.

The physical attachment of TARPs over AF injuries prevented degenerative remodeling across the IVD structure. Under static equilibrium, the swelling pressures in the NP create residual strains in the AF that exceed 10% strain in the outer AF (*33*). The release of this residual strain by annular lesions and herniation accelerate the degeneration of the surrounding tissue by instigating aberrant remodeling, short-term apoptosis, and the adoption of atypical fibrotic cellular phenotypes (*24*). In our model of annular injury, where the anterior AF was disrupted, unrepaired levels showed aberrant fibrotic remodeling, AF necrosis, and cellular apoptosis along the posterior AF after 4 weeks. These aberrant changes were not present in cervical discs that received TARP implantation, indicating that the mechanical reinforcement of the AF provided by the repair scaffolds may have helped to reestablish residual strains and prevent disc-wide remodeling. This promising finding is important for the long-term maintenance of disc health.

The delivery of Anakinra through TARP implantation also provided several benefits with potential to prolong disc health after injury. Some of the main hallmarks of disc degeneration include the depletion of NP proteoglycan content and a loss of the NP/AF boundary (*34–36*). Compared to empty TARP groups, Anakinra-loaded TARPs demonstrated an improved retention of NP proteoglycan content and the NP/AF boundary. This was accompanied by an increased infiltration of scar tissue along the anterior AF, which provided larger closure of the annular lesion and prevented NP displacement through the annular tear. This deep scar infiltration contrasts the weak scar deposition observed in human specimens and animal models of disc herniation where scar infiltration is limited to the outer third of full thickness annular lesions (*8, 37–39*). Anakinra provision also decreased osteolysis that was apparent with empty TARP delivery, indicating that blocking IL-1β signaling helped to attenuate the body’s response to the foreign material. Therefore, blockade of IL-1β signaling local to the injury site represents a promising therapeutic target to supplement annular closure strategies for disc herniation management.

The TARP annular closure system shows great potential for further preclinical and clinical development, yet this study is not without limitations. Although our study was limited in sample size and only assessed the short-term one-month timepoint, clear signs of therapeutic benefits are apparent. Future long-term assessments will determine if these early signs of improved repair result in the retention of disc health over longer study durations. Our injury model was limited to the anterior AF due to ease of surgical access, yet investigation of the effects of TARPs for the repair of posterolateral AF injuries would better recreate the herniation injuries mostly commonly observed in the clinic. This requires further development of an implantation strategy that is minimally invasive and provides safe access to the site. Additional investigations exploring the delivery of other molecules that could aid in annular healing, such as anti-apoptotic drugs and pro-anabolic agents, which may supplement our current approach for AF repair. Optimization of the use of distinct MAMC populations to enable sequential delivery of multiple agents may also be necessary to address the multiple phases of dense connective tissue repair. Altogether, the findings from this study support continued development of the TARP system and demonstrate that simultaneous repair and provision of molecules to the AF for the treatment of disc herniations is feasible and promising for clinical translation.

## MATERIALS AND METHODS

### Study Design

The objectives of this study were to develop tension-activated repair patches (TARPs) containing mechanically-activated microcapsules (MAMCs) and to elucidate the reparative effects of TARP-mediated annular repair and anti-inflammatory drug provision in vivo in a goat cervical disc injury model. We hypothesized that TARPs would provide structural reinforcement at the site of injury and enable anti-inflammatory drug delivery through MAMC mechano-activation, attenuating inflammation-induced matrix degradation.

TARPs were fabricated with different rhombus stamping patterns through variations in the longest diagonal of the rhombi (δ). The effects of stamp δ on MAMC patency after melt stamping was investigated (n=4/type), followed by characterization of TARP biomechanical properties (n=5/type). Local strain transfer under tension in response to stamp δ was investigated (n=5/type) and the effects of these differences on MAMC mechano-activation under increasing cycles of dynamic tensile loading at physiologic strains was investigated (n=4/type). To probe the effects of stamp design on fiber alignment and cellular morphology, fiber alignment and dispersity were measured post-stamping (n=4/type, n=4 ROIs/scaffold) and the morphology of human AF cells seeded on TARPs of different δ was assessed (n=4/type, n=30 ROIs/scaffold). To verify that MAMC fabrication did not alter inflammatory drug bioactivity, the interleukin 1 receptor antagonist drug, Anakinra, was encapsulated in MAMCs and its effect before and after encapsulation on the expression of catabolic genes by human AF cells, cultured in the presence of IL-1β, was assessed (n=4/group). Furthermore, a concentration sweep using different concentrations of Anakinra was performed to determine whether high drug concentrations affected AF cell viability (n=5/group). These studies enabled us to establish the most suitable stamping pattern for TARP *in vivo* mechano-activation upon delivery to injured discs.

To test the regenerative potential of TARP-mediated repair and Anakinra delivery, TARPs were tested in the goat cervical spine. Our *in vivo* study was approved by the University of Pennsylvania Institutional Animal Care and Use Committee (IACUC) and all surgeries followed the guidelines recommended by the committee. A total of 8 animals underwent surgery, consisting of annular injury of both C2-3 and C3-4 cervical discs followed by TARP repair of one level. Of the animals, n=4 received empty TARP repair (E-scaffold) and n=4 received Anakinra-loaded TARP repair (A-scaffold). Adjacent C4-5 levels were used as uninjured controls. Animals were sacrificed after 4 weeks, after which all motion segments underwent MRI T2 mapping, biomechanical testing, micro computed tomography imaging, and histological analysis. Due to level-to-level variations in MRI T2 relaxation times, level-matched uninjured controls from n=6 age-matched goats were used for MRI T2 map comparisons. One animal that received the A-scaffold was excluded from the study due to anatomical abnormalities found throughout the cervical spine. For each data set, outliers were identified as being outside 1.5x the interquartile range in Tukey box plots and were removed from the set before group comparisons.

## Funding

This work was supported by the United States Department of Veterans Affairs (I21 RX003447, IK6 RX003416, and IK2 RX003118) and the National Institutes of Health (P30 AR069619 and R01 AR071340).

## Author contributions

Conceptualization: APP, SEG, HES, RLM

Methodology: APP, SEG, EDB, MWH, TPS, HES, RLM

Investigation: APP, SEG, CSF, BSO, EDB, RLH, HMZ, MWH, TPS, HES, RLM

Visualization: APP, SEG, BSO

Funding acquisition: SEG, HES, RLM

Project administration: APP, SEG, HES, RLM

Supervision: SEG, GRD, DL, MWH, TPS, HES, RLM

Writing – original draft: APP, SEG, RLM

Writing – review & editing: All authors

## Competing Interests

Authors declare that they have no competing interests. A U.S. provisional patent for this work was filed titled “A Mechano-Responsive Nanofibrous Patch for the Delivery of Biologics in Load-Bearing Tissues”, No. 63/306,647. Co-inventors include APP, SEG, HES, and RLM.

## Data and Materials Availability

All data are available in the main text or the supplementary materials.

## SUPPLEMENTARY MATERIALS

### Materials and Methods

#### Nanofibrous Scaffold Fabrication

Dual component nanofibrous scaffolds composed of poly(e-caprolactone (PCL) (Shenzhn Esun Industrial Co., PCL 800C) and 200 kDa poly(ethylene oxide) (PEO) (Polysciences, 17503) were fabricated via electrospinning as previously described (*15, 16*). Briefly, a 14.3% w/v solution of PCL dissolved in a 1:1 mixture of tetrahydrofuran and N,N-dimethylformamide was made. Separately, PEO was dissolved in 90% EtOH to yield a 10% w/v solution. 50:50 PCL/PEO sheets (300-350 μm thick) were fabricated by simultaneously collecting PCL and PEO onto a grounded, rotating mandrel to form aligned nanofibers (**Fig. S1A**).

PCL-PEO scaffold strips were cut to required dimensions, with the longest length along the direction of fiber alignment. Scaffolds were hydrated through a gradient of EtOH (100%, 70%, 50%, 30%, and 2X phosphate buffered solution (PBS). For experiments requiring sterility, sterile washes were performed under a tissue culture hood. Note that hydration of scaffolds removed the PEO fraction (*15, 16*).

#### Fabrication of Mechanically-Activated Microcapsules

MAMCs were fabricated through the generation of water-in-oil-in-water double emulsions utilizing three liquid phases flowed through a glass capillary microfluidic device as previously established (**Fig. 1A**) (*18, 40*). The inner phase was composed of a pH 7.4 aqueous solution containing 1 mg/mL bovine serum albumin (BSA) as a model drug or 2 mg/mL of purified interleukin-1 receptor antagonist (IL-1ra) known as Anakinra (Sobi, Kineret^TM^), with 0.01% w/v AlexaFluor488-conjugated BSA (Invitrogen, A13100) for fluorescent visualization. Anakinra was purified from its base suspension solution using a 10 kDa filter (Millipore, UFC5010124), after which it was resuspended in PBS (pH7.4) containing AlexaFluor488-conjugated BSA to the desired concentration. Solution concentration was determined using the Pierce^TM^ BCA Protein Assay (ThermoFisher, 23250). The middle phase was composed of 85:15 lactide:glycolide poly(D,L-lactide-co-glycolide) (PLGA) copolymer (Lactel, B6006-1), dissolved in chloroform and fluorescently labeled with 100 mg/mL of Nile Red (Sigma, N3013). The outer phase was composed of 2% w/v poly(vinyl alcohol) in water. Double emulsions were collected in a pH 12 0.1% BSA in PBS solution with 0.95M of NaCl and left to harden over 72 hrs., during which time the chloroform evaporated from the middle phase, leaving behind a solidified PLGA wall. After that time, MAMCs were collected, washed with PBS, and stored at 4°C.

Fabrication efficiency and MAMC dimensions were assessed via confocal microscopy (Nikon A1R+), at 4X and 60X magnification, respectively. % Full was calculated by dividing full MAMCs over total MAMCs (n=5 counts/MAMC batch of >1000 MAMCs/count). MAMC dimensions were calculated using ImageJ, with n=20 images/batch measured.

#### Tension-Activated Annular Repair Patch Fabrication

TARPs were fabricated by melt-stamping MAMCs in between two hydrated PCL-PEO scaffold strips cut to dimensions required for each experiment (**Fig. 1B**). For assessment of stamping effect on MAMC patency, mechanical testing, local optical strain tracking, and fiber alignment/cellular morphology assessment, scaffold strips measuring 30 mm in length and 10 mm in width were used. For dynamic tensile loading, scaffold strips were cut to 60 mm in length and 10 mm in width. Finally, for in vivo studies, scaffold strips measuring 10 mm in length and 3.5 mm in width were used (**Fig. 1C**).

To melt-stamp scaffolds, 3D-printed metal stamps displaying a rhombus pattern with rhombus geometries that varied in longest rhombus diagonal length (δ) were used. Stamps with pattern δ = 3, 4, and 5 mm were utilized (**Fig. S1B**). A large stamp (l: 20 mm, w: 10 mm) was used to create all TARPs except for those used in the *in vivo* study (**Fig. S1B**), which were assembled using a smaller stamp (l: 10 mm, w: 3.5 mm) displaying the δ = 4 mm pattern. Stamps were heated at 80°C on a heat plate. Hydrated scaffold strips were laid flat on a flat surface and excess PBS was aspirated carefully. MAMCs, resuspended at the concentration of interest in a volume of 1.5 mL PBS/mm^2^ of scaffold (previously determined to not cause overflow during melt-stamping), were carefully pipetted over the flat scaffold strip. For all in vitro studies, TARPs were fabricated using a MAMC concentration of 700 MAMCs/mL. For in vivo implantation, empty TARPs (E-scaffold) were fabricated using 700 BSA-loaded MAMCs/mL while Anakinra-loaded TARPs (A-scaffold) were fabricated using 1,400 Anakinra-loaded MAMCs/mL to deliver 1 mg of Anakinra/TARP. After pipetting MAMCs onto the first scaffold strip, a second scaffold layer was carefully placed over the first, after removal of excess PBS, to cover MAMCs and the heated stamps were pressed over the scaffold-MAMCs-scaffold assembly for 3 seconds. Melt-stamped scaffolds were carefully detached from the stamp using tweezers and stored in PBS at 4°C.

#### Assessment of Stamping Effect on MAMC Patency

To assess the effects of stamping on MAMC patency, TARPs scaffold layers were carefully peeled apart to enable visualization of encapsulated MAMCs. MAMC inner solution and outer shell were fluorescently imaged using a ZEISS Axio Zoom V16 and n=20 regions of interest/scaffold were visualized for n=4 TARPs/stamp pattern. % Full was quantified as described above.

#### Mechanical Testing and Local Optical Strain Tracking

Before mechanical testing, the cross-sectional area of TARPs was measured using a custom-built laser device (*41*). Unstamped PEO-PCL scaffolds of comparable thickness were used as controls (thickness: ∼650 μm). TARPs were marked with black paint at the vertices of the rhombus patterns to enable local optical strain tracking of each rhombus during tensioning (**Fig. 2A**). Similarly, the unstamped control scaffolds were marked using a rhombus pattern. Small sections of sandpaper were superglued to both sides of each scaffold end (5 mm each end) to prevent slipping during testing. Scaffolds (n=5/type) were loaded in a universal test frame (Instron 5542, Instron, Norwood, MA) using custom grips at both ends and a 100N load cell. Loads were applied along the primary/longitudinal axis of the fibers. Specifically, a 60 sec. preload of 0.5N was applied, followed by 10 cycles of pre-conditioning at 0.5% strain at 0.1Hz and ramp to failure at 0.1% strain/second. During loading, images of the marked scaffolds were captured with a high-definition camera at 1 frame/sec (**Fig. 2A**).

A custom post-processing script (Matlab 2021a, Mathworks, Natick, MA) was developed to calculate mechanical properties based on “grip-to-grip” displacements and optical strain measures. Optical strain measurements of the rhombus-shaped patterns were achieved by identifying the 4 nodes of a given rhombus and tracking the change in position of the centroid of each node on every image. Tracked nodes were used to calculate the local axial (ε_y_) and transverse (ε_x_) strains over the course of the mechanical test. Slopes of the linear regions of force-displacement were used to derive values for stiffness.

#### Dynamic Mechanical Loading of TARPs

Using a custom-built bioreactor, n=4 TARPs/stamping pattern were dynamically tensile loaded at a time in a PBS bath to 6% strain at 1Hz, for 300, 1,800, or 3,600 cycles (*42*). The effects of loading cycles on MAMC patency were visualized using an AxioZoom and quantified as described above. Areas in which the stamps melted the scaffolds and MAMCs upon contact were not included in the quantification.

#### Fiber Alignment and Cell Morphology Changes with Stamp Geometries

##### Fiber Alignment and Dispersity Measurements

To assess fiber alignment changes with melt stamping, fiber autofluorescence was captured for unstamped PCL-PEO scaffolds and TARPs made with different stamp geometries (n=4 scaffolds/type, with n=4 ROIs/scaffold) under the DAPI channel using fluorescent confocal microscopy (Nikon A1R+, 20X magnification). Fiber alignment offset from 90° and alignment dispersity was calculated using the Directionality plugin in ImageJ. Fiber alignment maps overlayed over analyzed images were generated with the same plugin.

##### Cell Alignment and Aspect Ratio Analysis

The effects of melt stamping and stamp geometries on AF cell morphology and organization were assessed through visualization and quantification of cellular alignment and aspect ratio. Human AF cells were cultured on fibronectin-coated unstamped scaffolds and TARPs with different stamp geometries (n=4/type) at cells/mm^2^. After 6 days of culture, cells were washed 3x with PBS, after which they were fixed with 4% paraformaldehyde for 20 mins. at room temperature (RT). After fixation, the cells were washed 3x with PBS. Blocking was performed by adding blocking solution composed of 3% BSA in PBS for 1 hr. at RT. After blocking, cells were washed with PBS 3x, and cells were then stained with Draq5 (ThermoFisher, 62251) nuclear stain (1:1000) and Alexa Fluor 488 Phalloidin (ThermoFisher, A12379) (1:500) for 1 hr. at RT. After staining, cells were rinsed with PBS 2x and fluorescent confocal imaging was performed (Nikon A1R+, 20X magnification).

Cell aspect ratio and alignment angle were measured using CellProfiler, ver 3.1.9. Nuclei were viewed in n=30 ROIs/scaffold using the Draq5 signal and were segmented based on a minimum cross entropy algorithm with nuclei excluded below 5 and above 20 pixels. Nuclei were also excluded that were touching the image boundary. Cells were identified via propagation from the identified nuclei and segmented using a minimum cross entropy algorithm. Major and minor axes of the cells were measured, and their ratio (long to short axis) was presented as cell aspect ratio. The angle of the long axis of the cell was recorded to describe cell orientation compared to the scaffold fiber orientation.

#### Bioactivity Assessment of MAMC-Encapsulated Anakinra

##### In Vitro Culture and Treatment of Cells

To determine if the MAMC fabrication process affected the bioactivity of Anakinra after encapsulation, the effects of MAMC-encapsulated Anakinra were compared to those elicited by soluble Anakinra (**Fig. S3**). For this, human AF cells (Articular Engineering) (passage 2) were seeded at 6,000 cells/cm^2^ in 12-well plates with basal media composed of DMEM supplemented with 10% fetal bovine serum and 1% Penicillin-Streptomycin-Fungizone. After 4 days (∼70% confluence), the cell media was removed, and cells received basal media with 10 ng/mL of interleukin 1β (IL-1β) (R&D Systems, 201/LB/CF). Media was additionally supplemented with soluble Anakinra or the contents of Anakinra-encapsulating MAMCs at 0, 10, 100, 500 or 1,000 ng/mL, with n=4 wells/group receiving each treatment. MAMC contents were extracted by crushing MAMCs using a pestle. Anakinra concentrations were measured from extracted contents using the Pierce^TM^ BCA Protein Assay (ThermoFisher, 23250). Negative controls that did not receive any kind of IL-1β or Anakinra treatment were included. Media was replenished every 3 days, including all supplementing factors for each treatment group. After 6 days of treatment, cells were trypsinized, pelleted, and frozen.

##### Gene Expression Analysis via Quantitative Real Time RT-PCR

RNA isolation of digested samples was performed using the Directzol RNA Miniprep kit with DNAse-I treatment to remove trace DNA before RNA elution (Zymo Research, R2050). RNA was quantified via Nanodrop spectrophotometry. cDNA was synthesized using the SuperScript™ IV VILO Master Mix (Invitrogen, 11756050) according to the manufacturer’s protocol. Relative quantitative RT-PCR was run using Fast SYBR™ Green Master Mix (ThermoFisher, 4385618) for 40 cycles with validated primers (**Table S1**). Changes in gene expression were reported as 2^-DDCT^. ΔC_T_ for a given sample and gene of interest was calculated by subtracting the C_T_ value for the housekeeping gene (GAPDH) from the C_T_ value for the gene of interest. For a given gene, ΔΔC_T_ for a sample was calculated by subtracting average ΔC_T_ for the untreated negative control samples from the ΔC_T_ calculated for the sample.

##### Effects of Anakinra on Cell Viability

To test the effects of different Anakinra concentrations on cellular viability, human AF cells were cultured for 3 days in basal media, after which 0, 10, 100, 500 or 1,000 ng/mL of soluble Anakinra was added directly into the media, with n=5 wells/treatment. Media was replenished, including all supplementing factors for each treatment group, every 3 days. After 6 days, the media was removed and fresh media with alamarBlue^TM^ Cell Viability Reagent (ThermoFisher, DAL1025) was added, following manufacturer’s instructions. Empty wells also received the mixture as a background reference. After 4 hrs. of incubation at 37°C, fluorescence was measured using a spectrophotometer. Fluorescent intensity was calculated by subtracting the empty well reference measurement from each well measurement.

#### *In Vivo* Assessment of TARPs Using a Goat Cervical Disc Annular Injury Model

##### Injury Model & TARPs Implantation

Male (n=1) and female (n=7) skeletally mature (∼3 years of age), large frame goats were utilized. Under general anesthesia and using standard aseptic techniques, the animals underwent a surgical procedure at the C2-3 and C3-4 levels of the cervical spine to create an annular injury, with or without TARP implantation for repair of the induced injury (**Fig. 3A**). The ventral cervical spine was exposed using the anatomical plane between the trachea/esophagus and the carotid sheath. Intraoperative fluoroscopy was used to identify the vertebral bodies of C2 through C4. Soft tissues were bluntly and sharply dissected to expose the IVD of C2-3 and C3-4. Annular injury was induced, consisting of a cruciate partial thickness AF laceration (l: 5 mm, w: 2.5 mm, d: 4 mm), followed by full thickness puncture of the anterior AF through the middle of the X laceration (7mm depth) with a 14G needle (outer diameter: 2.1 mm) (**Fig. 3B-C**). A TARP (δ = 4 mm) loaded with BSA-encapsulating MAMCs (E-scaffold) or Anakinra-encapsulating MAMCs (A-scaffold) (l: 10 mm, w: 3.5 mm) was placed over the defect and sutured using 6-0 Gore-Tex monofilament suture (**Fig. 3D**). Sutures were placed in both corners of the cruciate annulotomy and the TARP was secured to each corner (**Fig. 3E**). Running sutures along the superior and inferior margins were used to further secure the TARP in place over the induced injury. The incision was then closed in layers and the animals recovered from anesthesia, after which animals were returned to standard housing consisting of 12 ft x 12 ft stalls. Animals were euthanized at 4 weeks post-implantation, and the cervical spines harvested en bloc and stored frozen at -20 °C until thawing for MRI scans and mechanical testing analyses.

##### Magnetic Resonance Imaging & Analysis

MRI scans of cervical spines were performed using a 3T scanner (Siemens Magnetom TrioTim). T2-weighted mid-sagittal images (5 mm slice thickness, 0.5 mm in plane resolution, TR/TE = 4,540/123 ms) were obtained. A series of images for T2 mapping (6 echoes, TE = 13 ms, 5 mm slice thickness, 0.5 mm in plane resolution) were also obtained.

Average T2 maps for each experimental group were generated using a custom MATLAB code, as previously described (*20*). Due to level-to-level variations in AF and NP T2, C3-4 uninjured healthy control levels from goat cervical spines used in a separate study (n=6) were used for T2 comparisons. T2 measured along the central Y axis (y=0) of normalized T2 maps generated through the MATLAB code were plotted for each specimen to show changes in T2 signal along the central axis of each disc. Average AF and NP T2 values were obtained for each specimen by measuring rectangular and circular regions of interest for the AF and NP regions, respectively, on T2 maps using ImageJ.

##### Motion Segment Biomechanical Testing & Analysis

Motion segments (vertebra-disc-vertebra units) were prepared for compression testing by carefully dissecting musculature around the disc and removing posterior and lateral boney elements with a hand saw. Ink spots were placed on the vertebral bone immediately distal and proximal to the disc for optical tracking during testing (**Fig. S9A**). Motion segments were then potted in a low melting temperature indium casting alloy (McMaster-Carr) in custom fixtures and subjected to a compression and creep testing protocol (Instron 5948) in a PBS bath with protease inhibitors. The testing protocol consisted of 20 cycles of compression from 0 to -100 N (0 to 0.14 MPa) followed by creep testing consisting of a 1 second step load to -0.14MPa and a 60-minute hold.

A bi-linear fit of the 20^th^ compression curve was performed in MATLAB to quantify toe and linear region modulus, as well as maximum compressive strain for each sample. Creep displacement was calculated by fitting the creep test to a 5-parameter viscoelastic constitutive model in MATLAB, as previously described (*43, 44*).

##### MicroComputed Tomography Imaging & Analysis

To prepare motion segments for MicroComputed Tomography (μCT), samples were fixed for 7 days in formalin at 4°C. After fixation, samples were rinsed in PBS, wrapped in PBS-soaked gauze, and placed within the device for scanning. Motion segments were imaged at an isotropic 10 mm resolution using a Scanco Medical μCT50.

Cranial and caudal bony endplates (defined as the region between the intervertebral disc and growth plate) were manually contoured (**Fig. S4A**) and bone morphometry parameters, including bone volume over total volume (BV/TV), trabecular number (Tb. N.), and trabecular spacing (Tb. Sp) were calculated using the Scanco Medical Analyzer software. Regions where osteolysis was apparent on the cranial and caudal bony endplates were manually contoured and osteolysis volume was calculated. Scanco Medical Visualizer software was used to generate 3D reconstructions of specimen scans with color-coded display of bone density.

##### Histological Assessment of Disc Composition and Structure

Motion segments were decalcified (Formical-2000, Decal Chemical Corporation, Tallman, NY) and processed through paraffin. 10 μm sections around the mid-sagittal plane were collected. For all stains performed, sections from all experimental groups were stained simultaneously. Sections were co-stained for glycosaminoglycans and collagens using Alcian Blue and Picrosirius Red (AB/PSR), respectively. NP Alcian Blue staining intensity was measured using a circular region of interest in ImageJ. For the visualization of microscopic anatomy and cellular morphology, sections were stained with hematoxylin and eosin (H&E). To visualize collagen fibrils, elastin, bone, other hyaline structures, and cells, a one-step Mallory-Heidenhain (MH) stain was used (*45*) (**Fig. 6**). Collagenous scar infiltration area along the anterior and posterior AF was measured in the red channel using thresholds applied with ImageJ.

#### Schematics and Illustrations

All schematics and illustrations were generated using BioRender.com and/or Microsoft PowerPoint.

#### Statistical Analysis

Statistical analyses were performed using Prism 9 (Graph Pad Software Inc.), with significance defined as p-value < 0.05. Quantitative data is presented as mean ± standard deviation (s.d.) or mean ± standard error of the mean (SEM). The Shapiro-Wilk normality test was used to determine the need for non-parametric testing (alpha = 0.05). For each data set, outliers were identified as being outside 1.5x the interquartile range in Tukey box plots and were removed from the set before group comparisons. For comparisons between stamping patterns (MAMC patency post-stamping, bulk scaffold biomechanical properties, local strains, fiber alignment and dispersity, and cellular alignment and aspect ratio analyses), one-way ANOVA with Tukey’s multiple comparisons post-hoc analysis was employed. Similarly, for Alamar Blue and qPCR analyses for different concentrations of Anakinra, a one-way ANOVA with Tukey’s multiple comparisons post-hoc analysis was used. MAMC mechano-activation with dynamic tensile loading was analyzed using a two-way ANOVA with Tukey’s multiple comparisons post-hoc test. To characterize the effect of the induced annular injury, uninjured controls and injury groups were compared using an unpaired t-test with Welch’s correction. AF and NP T2 signal, disc biomechanical properties, bone morphological characteristics, anterior and posterior disc scar infiltration/remodeling area, and NP Alcian Blue intensity were analyzed using one-way ANOVA with Tukey’s multiple comparisons post-hoc analysis. Osteolysis measurements were compared using a non-parametric one-way ANOVA with Dunn’s multiple comparisons.

---

**Fig. S1.**
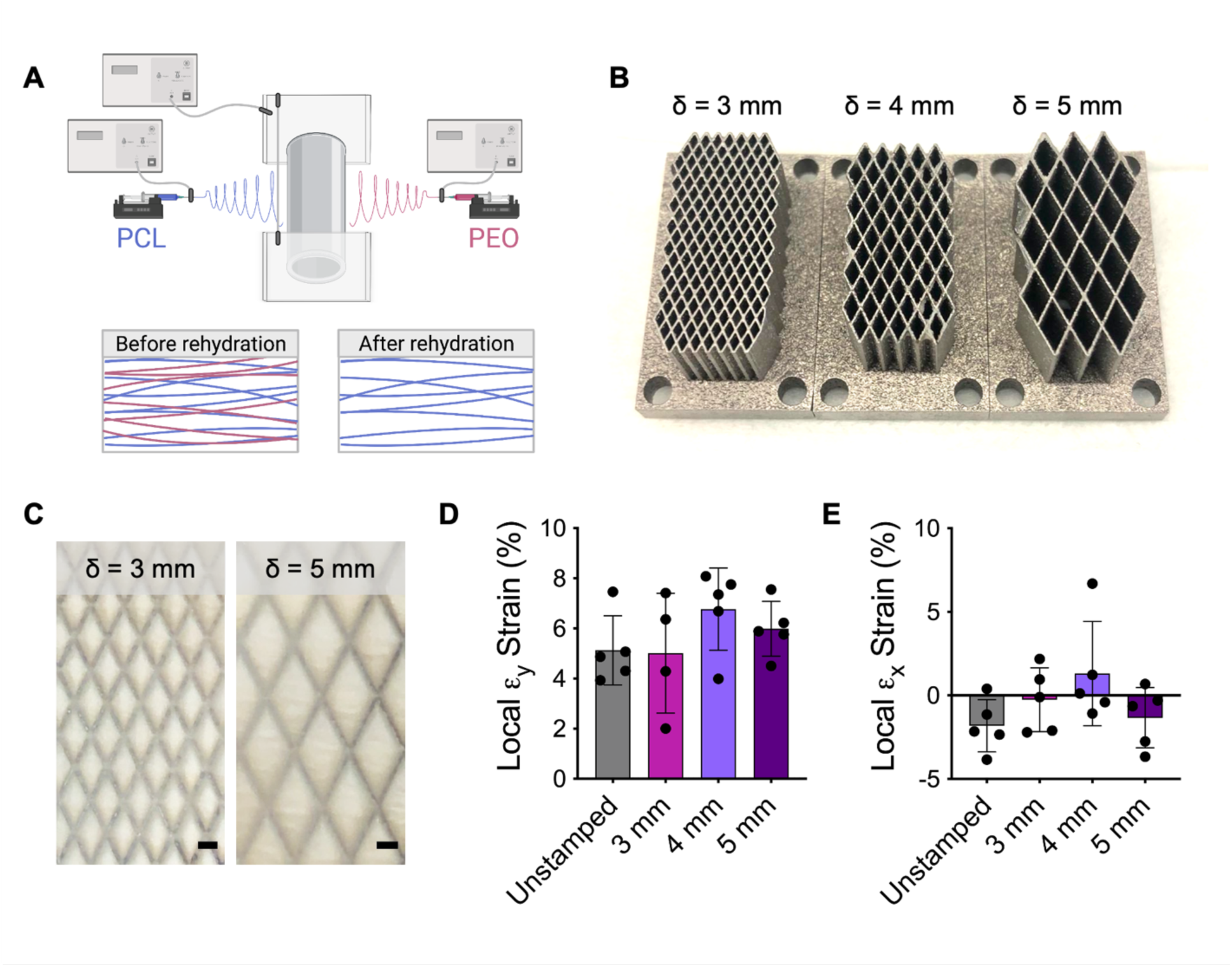
TARP fabrication and local strain measurements. (**A**) Schematic of fabrication of PCL-PEO nanofibrous scaffolds through simultaneous electrospinning and collection of fibers composed of each material on a rotating mandrel. (**B**) 3D-printed metal stamps displaying rhombus grids with varying rhombus geometries through the change in longest rhombus diagonal (δ), used for melt-stamping of TARPs. (**C**) Representative images of melt-stamped PCL-PEO layers with δ = 3 mm (left) and δ=5 mm (right) (scale: 1 mm). (**D**) Local longitudinal strain (ε_y_) and (**E**) transverse strain (ε_x_) (mean ± s.d.) in each rhombus at an applied Instron strain of 6% for unstamped scaffolds vs stamped groups (n=5/type).

**Fig. S2.**
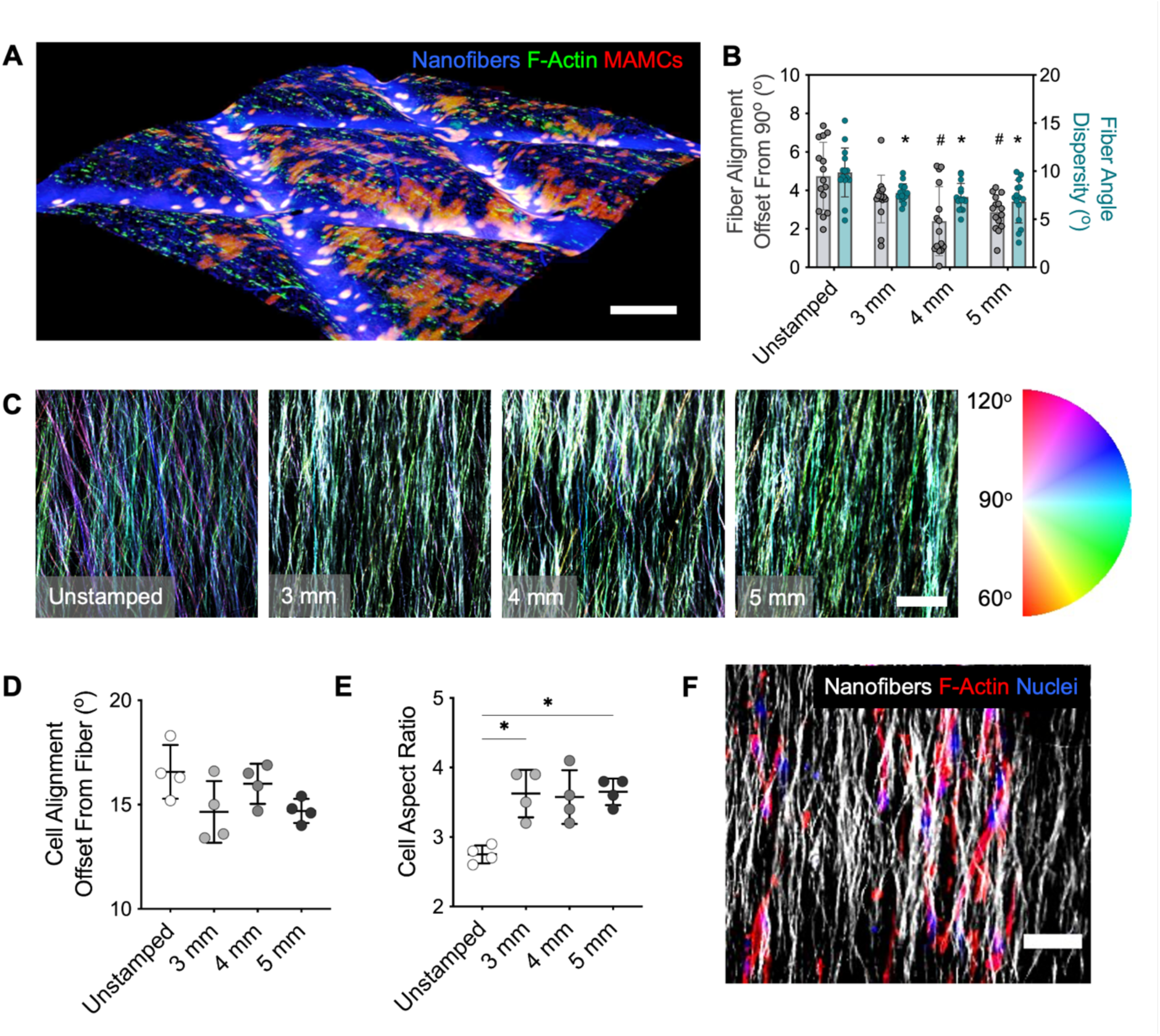
Effect of melt-stamping and rhombus geometry on fiber and cellular alignment. (**A**) 3D reconstruction of confocal z-stack for δ = 3 mm TARP seeded with human AF cells (blue: nanofibers, green: f-actin, red: MAMC shells) (scale: 500 μm). (**B**) Fiber alignment offset from 90° and fiber angle dispersity (mean ± s.d.) for unstamped and stamped scaffolds (n=4 scaffolds/type, with n=4 ROIs/scaffold). (**C**) Scaffold fiber orientation maps for unstamped and stamped TARPs with fibers colored based on orientation angle (scale: 50 μm). (**D**) Cell alignment offset from fiber direction and (**E**) cell aspect ratio (mean ± s.d.) for cells seeded on scaffolds (n=4 scaffolds/type, with n=4 ROIs/scaffold). (**F**) Representative confocal image of cells seeded on a δ = 4 mm TARP, with cell bodies (F-actin: red; nuclei: blue) oriented along the fiber direction (scale: 50 μm). Statistical significance denoted by # p-value < 0.05 vs. unstamped for fiber alignment offset, * p-value < 0.05 vs unstamped for fiber angle dispersity.

**Fig. S3.**
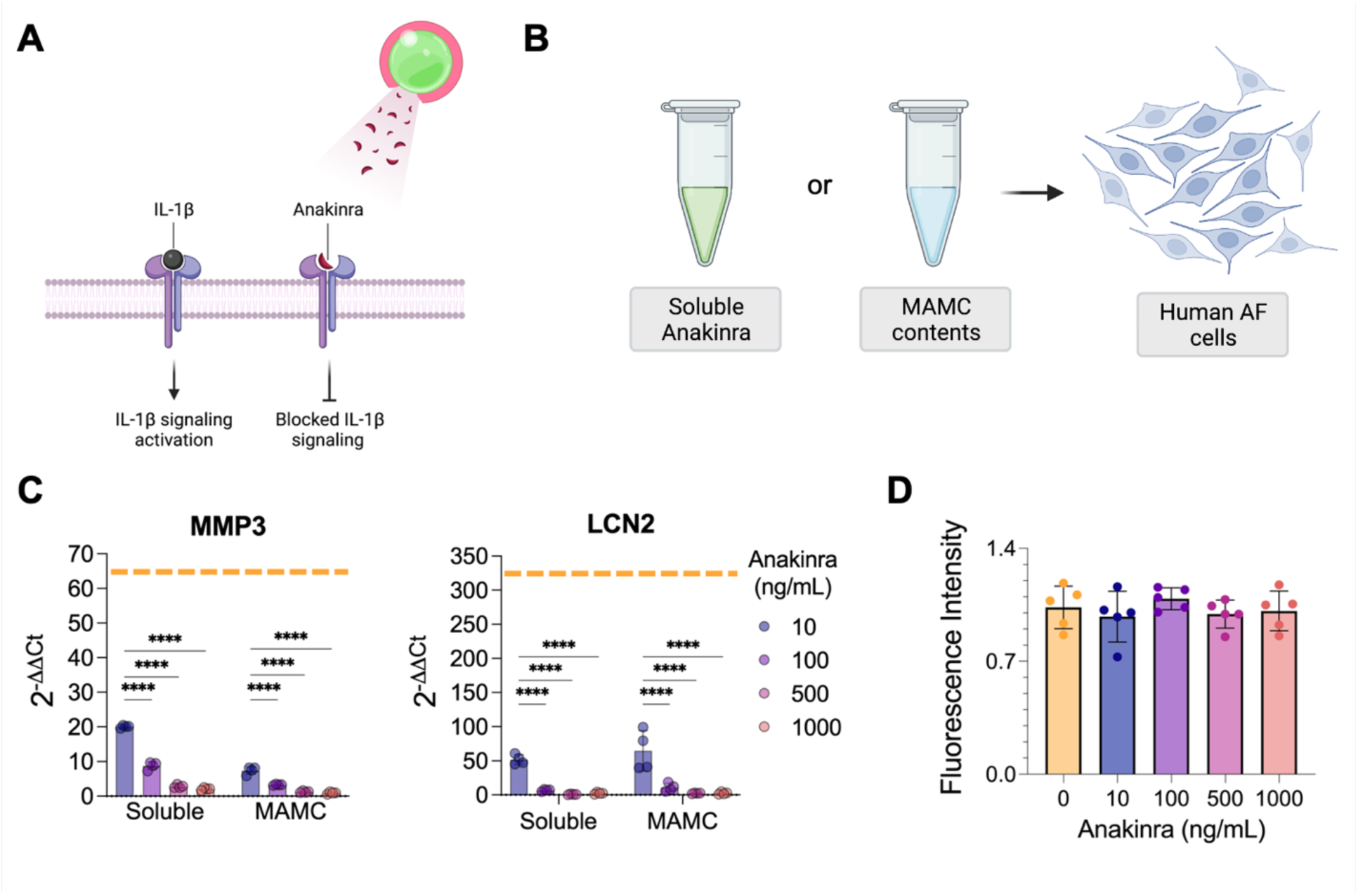
MAMC-encapsulated Anakinra retains bioactivity and potently blocks IL-1b-mediated effects on human AF cells. (**A**) Schematic of the mechanism of MAMC-mediated Anakinra delivery to block IL-1b signaling. (**B**) Schematic of the comparison between direct delivery of soluble Anakinra and the contents of Anakinra-loaded MAMCs on human AF cells cultured with or without IL-1b. (**C**) Alamar Blue fluorescent intensity (mean ± s.d.) of the media from AF cells cultured with different Anakinra concentrations for 6 days (n=5/treatment). (**D**) Relative gene expression (mean ± s.d.) for MMP3 (left) and LCN2 (right) for AF cells cultured in the presence of IL-1b with or without different doses of Anakinra delivered via soluble direct delivery or extracted from Anakinra-loaded MAMCs (n=4/treatment). Orange dashed line represents gene expression for cells that did not receive Anakinra. Statistical significance denoted by * p-value **** < 0.0001.

**Fig. S4.**
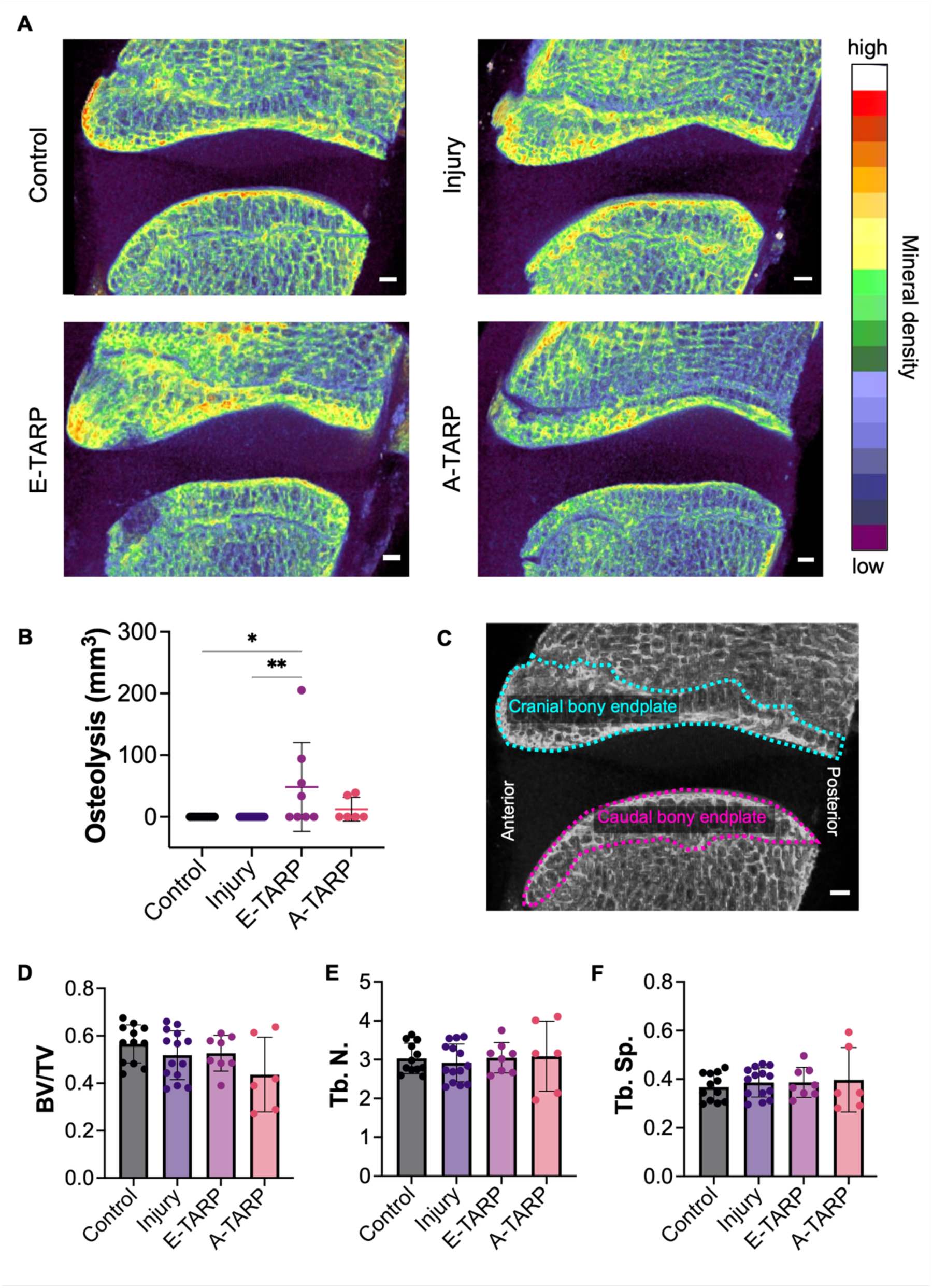
TARP-mediated Anakinra delivery attenuates implantation-mediated osteolysis in adjacent vertebras. (**A**) Sagittal view of representative bone density maps for motion segments in each group, showing osteolysis in the vertebras adjacent to the E-Scaffold-treated disc (scale: 1 mm). (**B**) Osteolysis volume measured across groups on the cranial and caudal bony endplates of the motion segments (mean ± s.d.). (**C**) Example contouring of cranial and caudal bony endplates along the sagittal plane that were morphed to created 3D contours for bone morphometric analysis in D-F (scale: 1 mm). (**D**) Bone volume to total volume ratio (BV/TV), (**E**) trabecular number (Tb. N.), and (**F**) trabecular spacing (Tb.Sp.) measured across groups on the cranial and caudal bony endplates of the motion segments (mean ± s.d.). Cranial and caudal endplates from the same animal are shown as individual data points. Statistical significance denoted by * p-value < 0.05, ** < 0.01.

**Figure S5.**
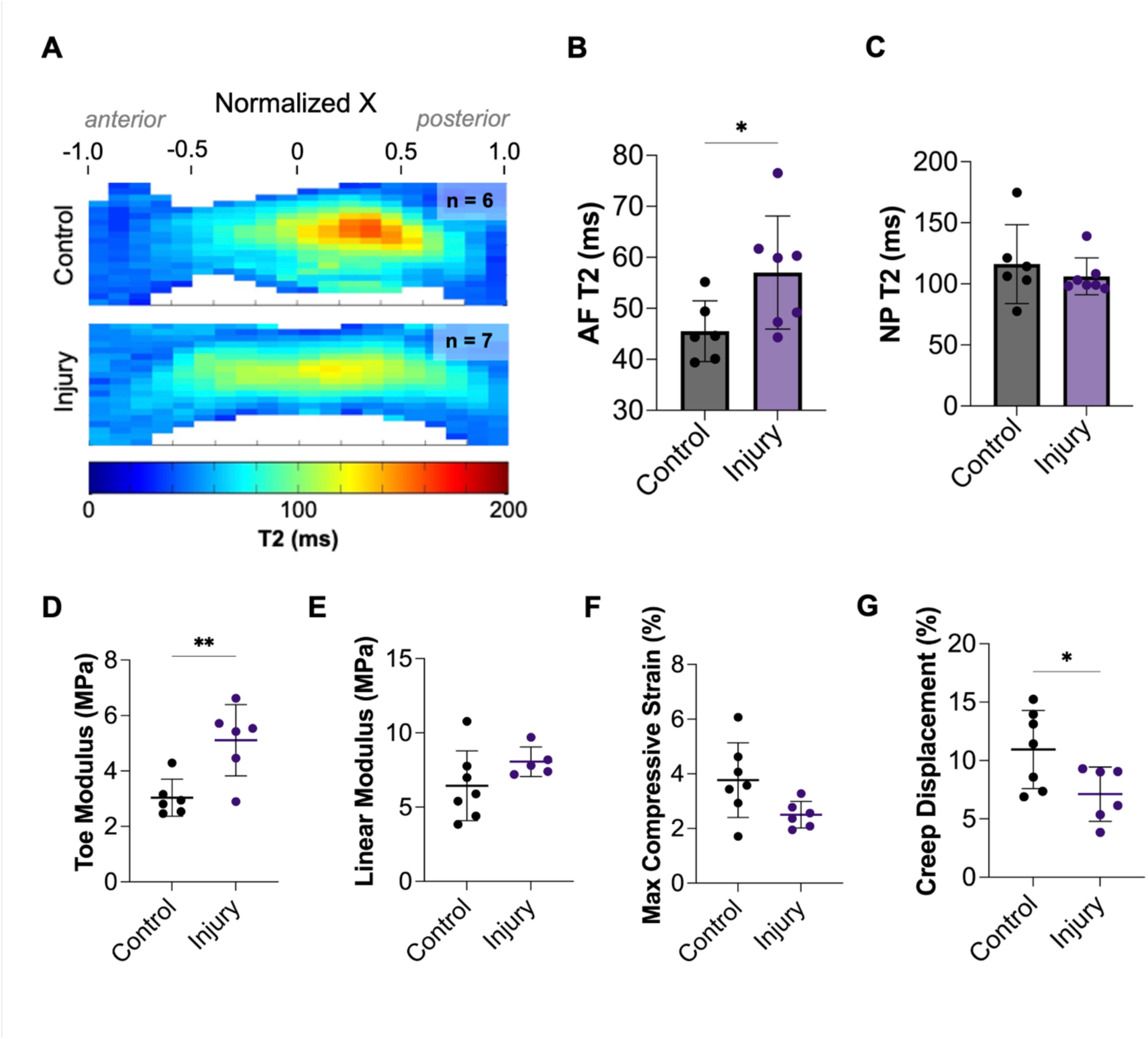
Effects of annular injury on T2 and biomechanics. (**A**) Average T2 maps for each group showing level-matched controls (n=6) and injury (n=7). (**B**) AF and (**C**) NP T2 measurements for rectangular and circular regions of interest, respectively, for uninjured vs injured discs (mean ± s.d.). (**D**) Toe modulus, (**E**) linear modulus, (**F**) maximum compressive strain, and (**G**) creep displacement (mean ± s.d.) for motion segments in each group (n=7 and n=6 for uninjured control and injured discs, respectively). Statistical significance denoted by * p-value < 0.05, ** < 0.01.

**Fig. S6.**
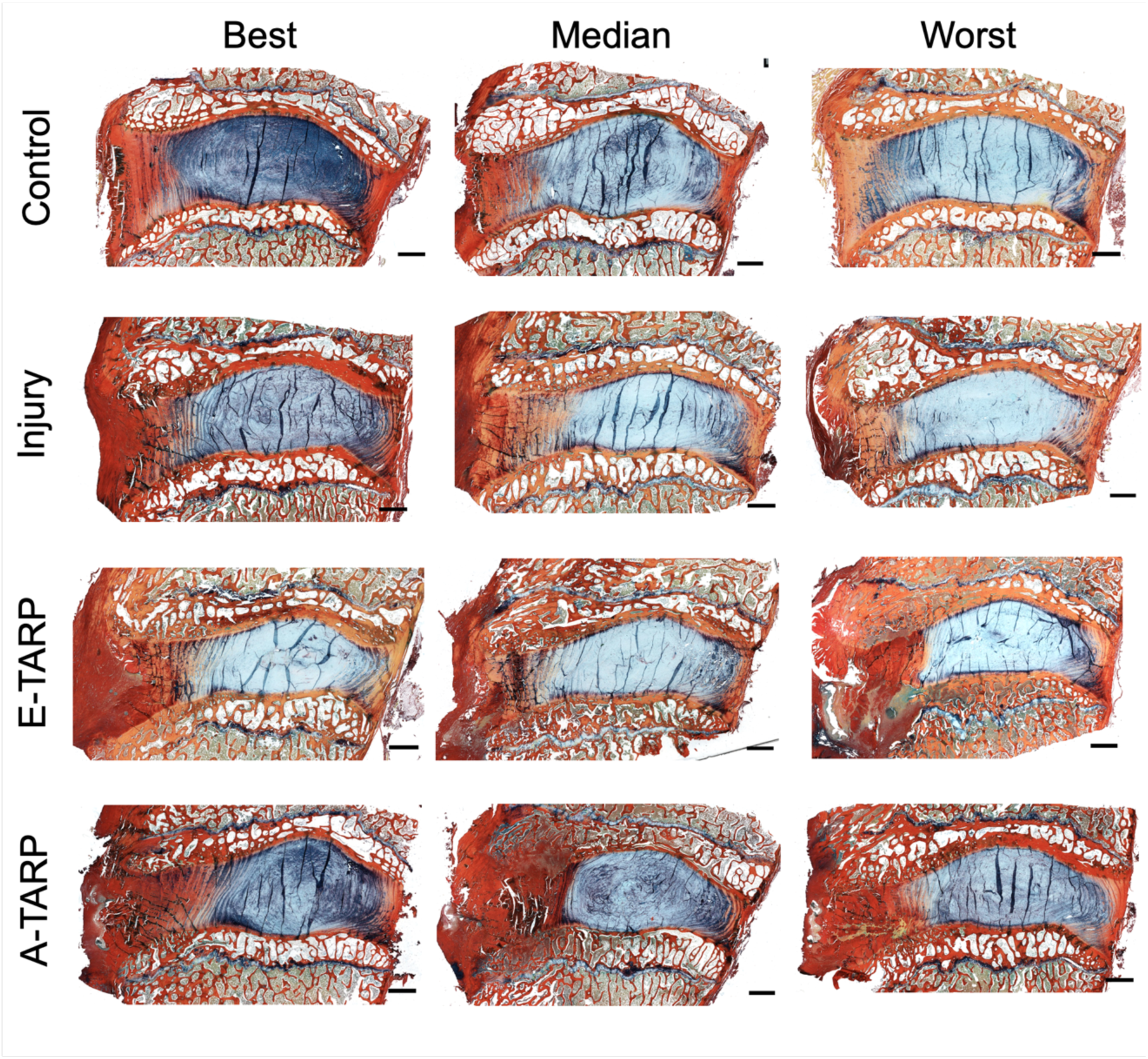
Biochemical compositional changes with annular injury and TARP repair. Alcian Blue-Picrosirius red-stained sagittal histological sections showing best, median, and worst specimens for each group, identified through NP Alcian Blue intensity measurements (scale: 2 mm).

**Fig. S7.**
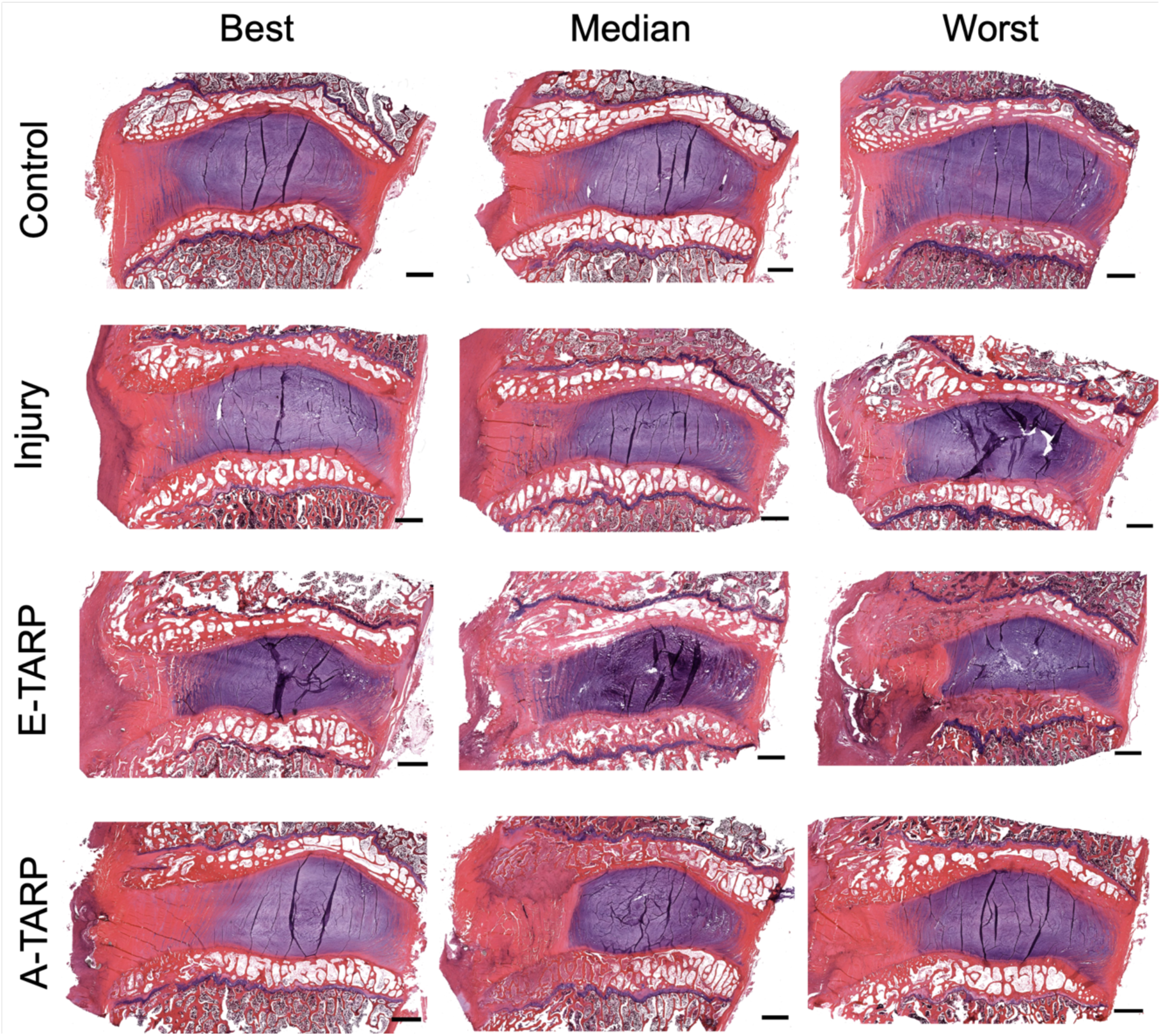
Structural and compositional changes with annular injury and TARP repair. H&E-stained sagittal histological sections showing best, median, and worst specimens for each group, identified through NP Alcian Blue intensity measurements (scale: 2 mm).

**Fig. S8.**
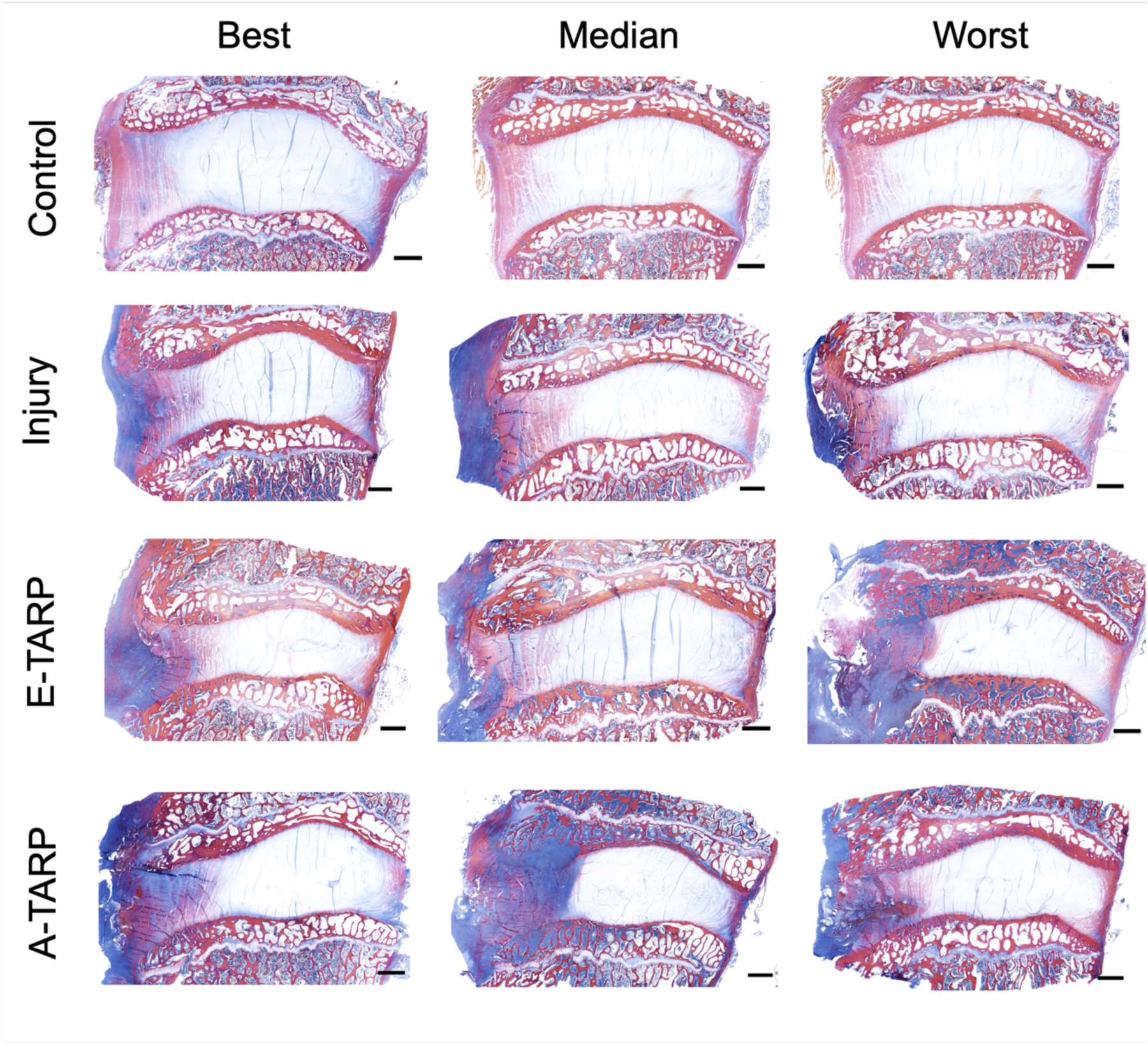
Collagenous scar infiltration and remodeling differences with injury and treatment. Mallory Heindenhain-stained sagittal histological sections showing best, median, and worst specimens for each group, identified through NP Alcian Blue intensity measurements (scale: 2 mm).

**Fig. S9.**
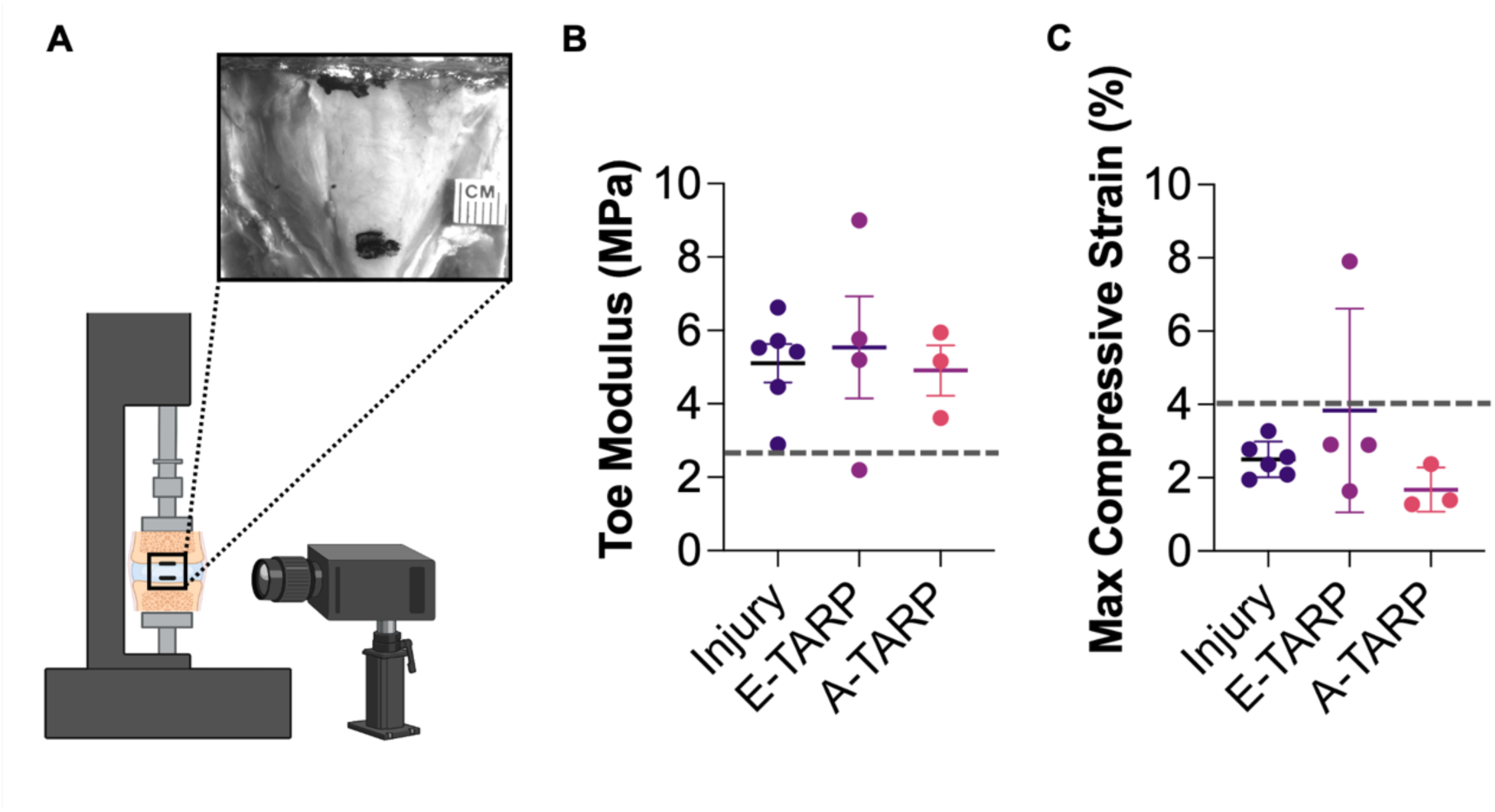
Injury-induced stiffening of the disc is not reversed by TARP-implantation. (**A**) Depiction of mechanical testing setup for motion segment compression and creep testing. PBS bath not depicted. (**B**) Toe modulus and (**C**) maximum compressive strain (mean ± s.d.) for motion segments in each group compared to uninjured controls (dashed line: control mean).

**Supplementary Table S1.**
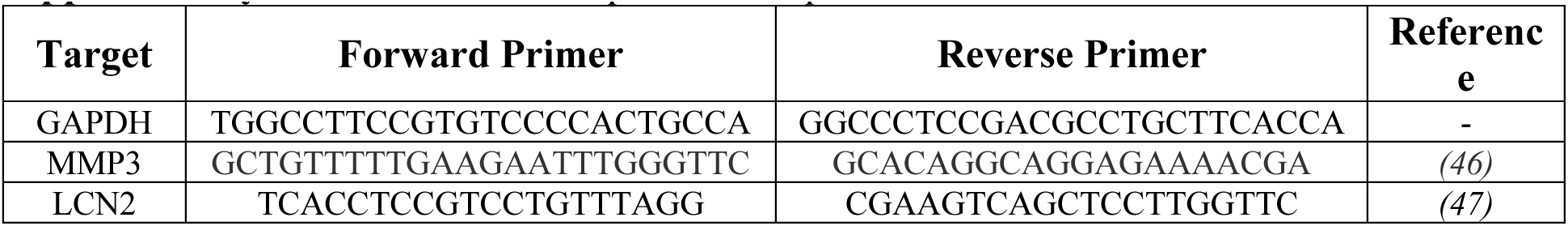
Primer sequences for quantitative RT-PCR.

## Notes

### Competing Interest Statement

The authors have declared no competing interest.

